# SNAI2-mediated direct repression of *BIM* protects rhabdomyosarcoma from ionizing radiation

**DOI:** 10.1101/2021.01.25.428112

**Authors:** Long Wang, Nicole R. Hensch, Kathryn Bondra, Prethish Sreenivas, Xiang Ru Zhao, Jiangfei Chen, Kunal Baxi, Rodrigo Moreno Campos, Berkley Gryder, Silvia Pomella, Rossella Rota, Eleanor Y. Chen, Javed Khan, Peter J. Houghton, Myron S. Ignatius

## Abstract

Ionizing radiation (IR) and chemotherapy are the mainstays of treatment for patients with rhabdomyosarcoma (RMS). Yet, the molecular mechanisms that underlie the success or failure of radiotherapy remain unclear. The transcriptional repressor SNAI2 was previously identified as a key regulator of IR sensitivity in normal and malignant stem cells through its repression of the proapoptotic BH3-only gene *PUMA.* Here, we demonstrate a clear correlation between SNAI2 expression levels and radiosensitivity across multiple RMS cell lines. Moreover, modulating *SNAI2* levels in RMS cells through its overexpression or knockdown can alter radiosensitivity *in vitro* and *in vivo.* SNAI2 expression reliably promotes overall cell growth and inhibits mitochondrial apoptosis following exposure to IR, with either variable or minimal effects on differentiation and senescence, respectively. Importantly, *SNAI2* knockdown results in a striking increase in expression of the proapoptotic BH3-only gene *BIM,* and ChIP-seq experiments establish that SNAI2 is a direct repressor of *BIM.* Since the P53 pathway is nonfunctional in the RMS cells used in this study, we have identified a new, P53-independent SNAI2/BIM axis that could potentially predict clinical responses to IR treatment and be exploited to improve RMS therapy.

**Highlights:**

- SNAI2 expression levels are directly correlated with protection from radiation in rhabdomyosarcoma.
- Loss of SNAI2 primes rhabdomyosarcomas for IR-induced apoptosis.
- SNAI2 directly represses the expression of the proapoptotic BH3-only gene *BIM.*

## Introduction

Ionizing radiation (IR), chemotherapy and surgery comprise the current standard of care for patients with rhabdomyosarcoma (RMS), a pediatric malignancy of the muscle, and lead to greater than 70% tumor-free survival in children with this disease^1–3^. However, the survival rate for patients with disease relapse remains dismal at less than 30%^1–4^. IR is used to treat both primary tumors and metastatic lesions in relapsed RMS patients^5^. Remarkably, these patients often receive a cumulative dose of 36 to 50.4 Gy^5^; yet, an understanding of the pathways that regulate the IR-induced DNA damage response in RMS tumors remains incomplete. In this study, we identify SNAI2 as a critical radioprotector of RMS tumor cells and define the pathways downstream of *SNAI2* signaling that regulate the response to IR in RMS.

SNAIL genes comprise a family of transcriptional repressors important for epithelial morphogenesis during development (e.g., the epithelial-mesenchymal transition) and for cell survival^6–8^. The role of *SNAI2* in protecting normal hematopoietic stem cells (HSCs) from IR-induced apoptosis is well established^9, 10^. In radiosensitive cells (e.g., the lymphoid lineage), exposure to IR causes DNA double-strand breaks that trigger the activation of P53. P53-mediated induction of the BH3-only gene *PUMA* then leads to mitochondrial apoptosis^11^. However, HSCs are uniquely protected from IR-induced apoptosis due to a concomitant P53-mediated induction of *SNAI2* in these cells, which directly represses the expression of *PUMA^10^.* Recent studies also implicate SNAI2 as a regulator of the DNA damage response in normal mammary stem cells^12^, and *SNAIL* family members *SNAI1* and *TWIST* have been shown to regulate the IR-induced DNA damage response in breast cancer cells through regulation of *ZEB1*^13^. These studies suggest that the SNAIL family may have widespread importance in regulating the response to IR.

Not surprisingly, adult cancers and relapse disease often present with mutations in or loss of *TP53*^14, 15^. In a subset of these tumors that are still radiosensitive, IR has been shown to induce cell death through P53-independent mechanisms involving cell cycle checkpoint proteins and alternative DNA damage response pathways (Reviewed in^16, 17^). Interestingly, childhood cancers including RMS often retain wild-type *TP53*^18–21^. Mutations in *TP53* account for <6% of RMS primary tumors, yet they can be acquired during relapse and are associated with a poor outcome^19–21^. Since RMS tumors harboring either wild-type or mutant P53 are sensitive to IR^19–21^, both P53-dependent and -independent mechanisms of IR-induced cell death appear to be active in this disease. Our analysis show that SNAI2 protects RMS tumors from IR both *in vitro* and *in vivo.* Using RMS cell lines that express varying levels of SNAI2 and a dysfunctional P53 pathway^22, 23^, we show that levels of SNAI2 establish the degree of protection of RMS cell lines from IR, regardless of *TP53* mutation status, and that the proapoptotic BH3-only gene *BIM* is directly repressed by SNAI2 to confer protection from radiation. Our results suggest that SNAI2 is a major player in the response to IR in RMS and represents a promising target for the radiosensitization of RMS tumors during IR therapy.

## Results

### SNAI2 expression directly correlates with radiosensitivity in RMS cells

To better understand the factors regulating sensitivity to IR in RMS tumors, we used a panel of 4 representative human RMS cell lines Rh18, RD, Rh41, and Rh30. These cell lines include both Embryonal RMS (ERMS) and Alveolar RMS (ARMS) subtypes in which *TP53* is both mutant and wild-type (Table 1). Using an imaging-based platform to test different doses of IR on cell number/confluence, we found that Rh18 cells are relatively more sensitive to IR, with RD and Rh41 showing moderate sensitivity, and Rh30 the most radioresistant (Figure 1A-D, Supplemental Figure 1A-B). Since *SNAI2* is known to protect cells from IR in other tumors^9, 10^, we analyzed SNAI2 expression levels across RMS cell lines. Interestingly, the expression of SNAI2 was correlated with the degree of protection from radiation, with Rh18 showing low SNAI2 expression levels, RD and Rh41 showing moderate levels and Rh30 expressing high levels of SNAI2 protein. In contrast, the expression of proteins known to be involved post IR including SNAI1, MDM2, ZEB1, CHEK1 and CHEK2 were not correlated with radiosensitivity (Figure 1E, Supplemental Figure 1E). Analysis of *SNAI2* expression across RMS tumor lines indicates that there is a trend toward higher SNAI2 expression in ERMS tumors compared to ARMS, but that expression is variable across tumors (Figure 1F). Finally, analysis of The Cancer Genome Atlas (TCGA) and St. Jude PeCan Data Portal for *SNAI2* expression showed that sarcomas, including RMS, are among the cancers that express the highest levels of *SNAI2* and have higher expression levels compared to control tissue (Figure 1F and Supplemental Figure 1C-D). These results suggest that SNAI2 may have a pro-tumorigenic function in RMS.

**Figure 1.**
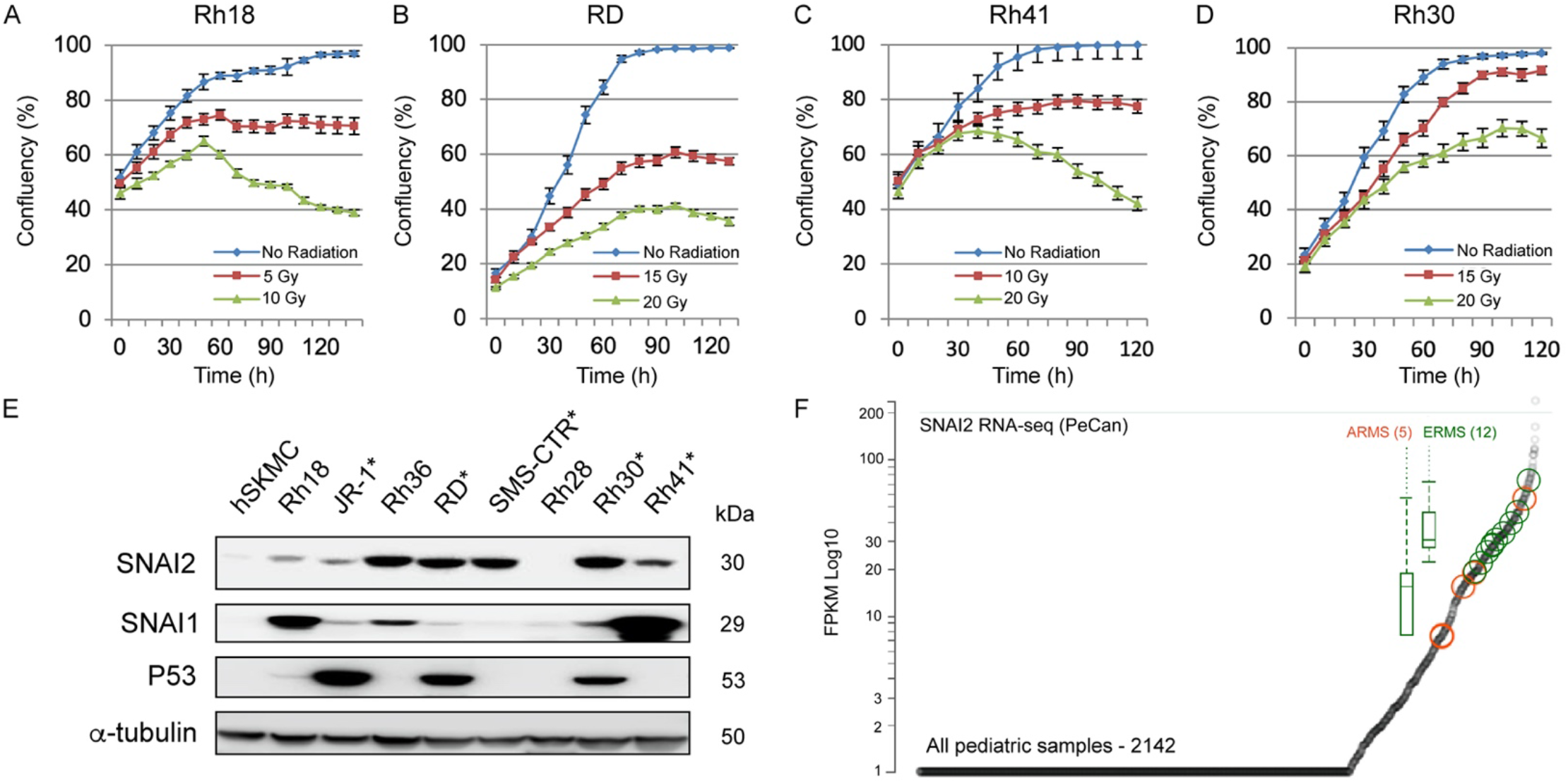
SNAI2 expression directly correlates with radiosensitivity in RMS cells. A-D. Rh18, RD, Rh41, and Rh30 cells were irradiated at 24h post imaging with varying levels of radiation and cell confluency (%) was assessed using Incucyte Zoom software based on phase-contrast images acquired from 0 h to 120 h. Error bars represent ± 1 SD. E. Western blot showing protein levels of SNAI1, SNAI2, and P53 in various parental RMS cell lines with skeletal human myoblast cells (hSKMCs) as a control. Asterisks (*) note RMS cell lines with known P53 mutations. F. PeCan SNAI2 RNA-seq data for ARMS (orange) and ERMS (green) tumors.

**Table 1.**
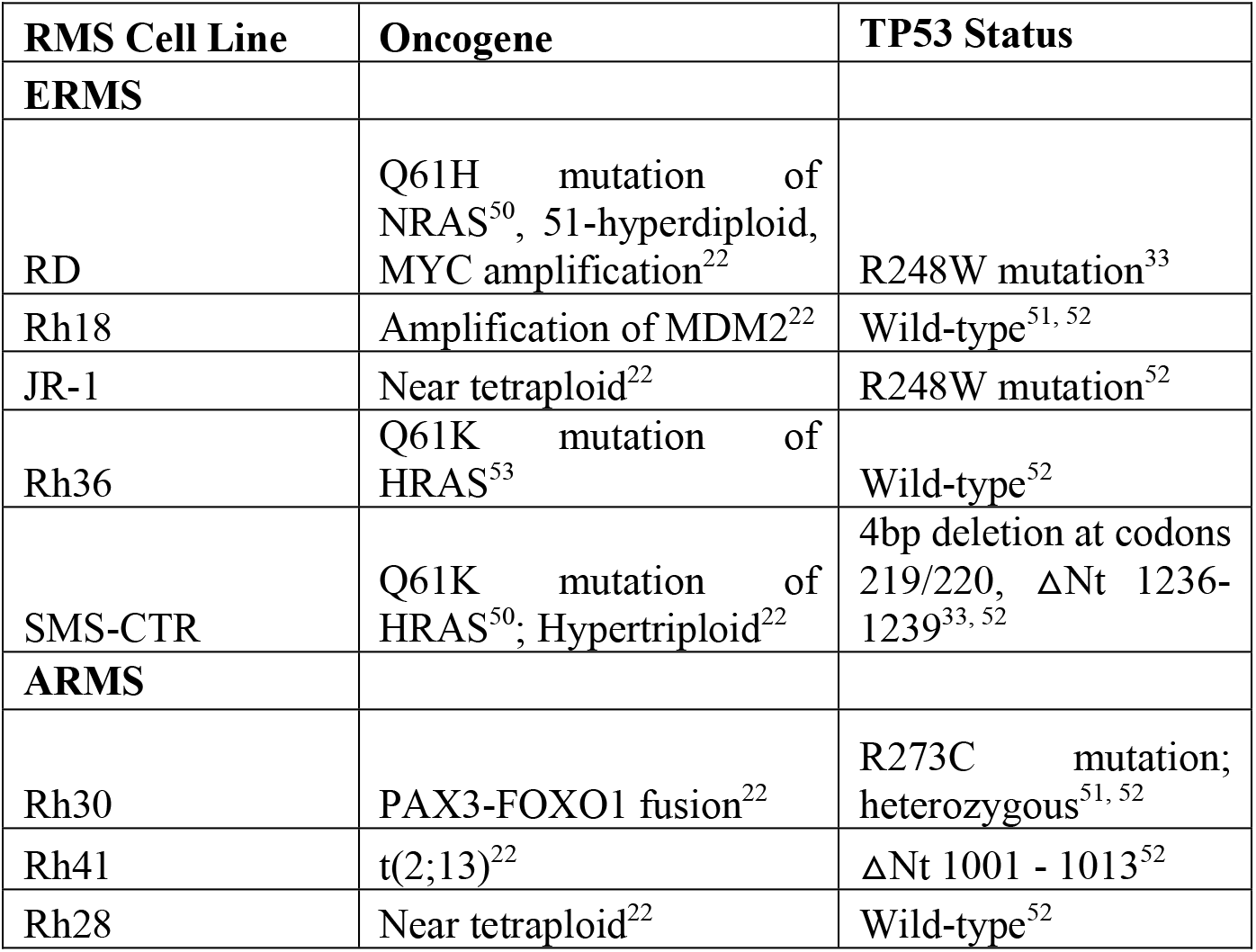

| RMS Cell Line | Oncogene | TP53 Status |
| --- | --- | --- |
| **ERMS** |  |  |
| RD | Q61H mutation of NRAS<sup>50</sup>, 51-hyperdiploid, MYC amplification<sup>22</sup> | R248W mutation<sup>33</sup> |
| Rh18 | Amplification of MDM2<sup>22</sup> | Wild-type<sup>51, 52</sup> |
| JR-1 | Near tetraploid<sup>22</sup> | R248W mutation<sup>52</sup> |
| Rh36 | Q61K mutation of HRAS<sup>53</sup> | Wild-type<sup>52</sup> |
| SMS-CTR | Q61K mutation of HRAS<sup>50</sup>; Hypertriploid<sup>22</sup> | 4bp deletion at codons 219/220, ΔNt 1236-1239<sup>33, 52</sup> |
| **ARMS** |  |  |
| Rh30 | PAX3-FOXO1 fusion<sup>22</sup> | R273C mutation; heterozygous<sup>51, 52</sup> |
| Rh41 | t(2;13)<sup>22</sup> | ΔNt 1001 - 1013<sup>52</sup> |
| Rh28 | Near tetraploid<sup>22</sup> | Wild-type<sup>52</sup> |

**Table 2.**
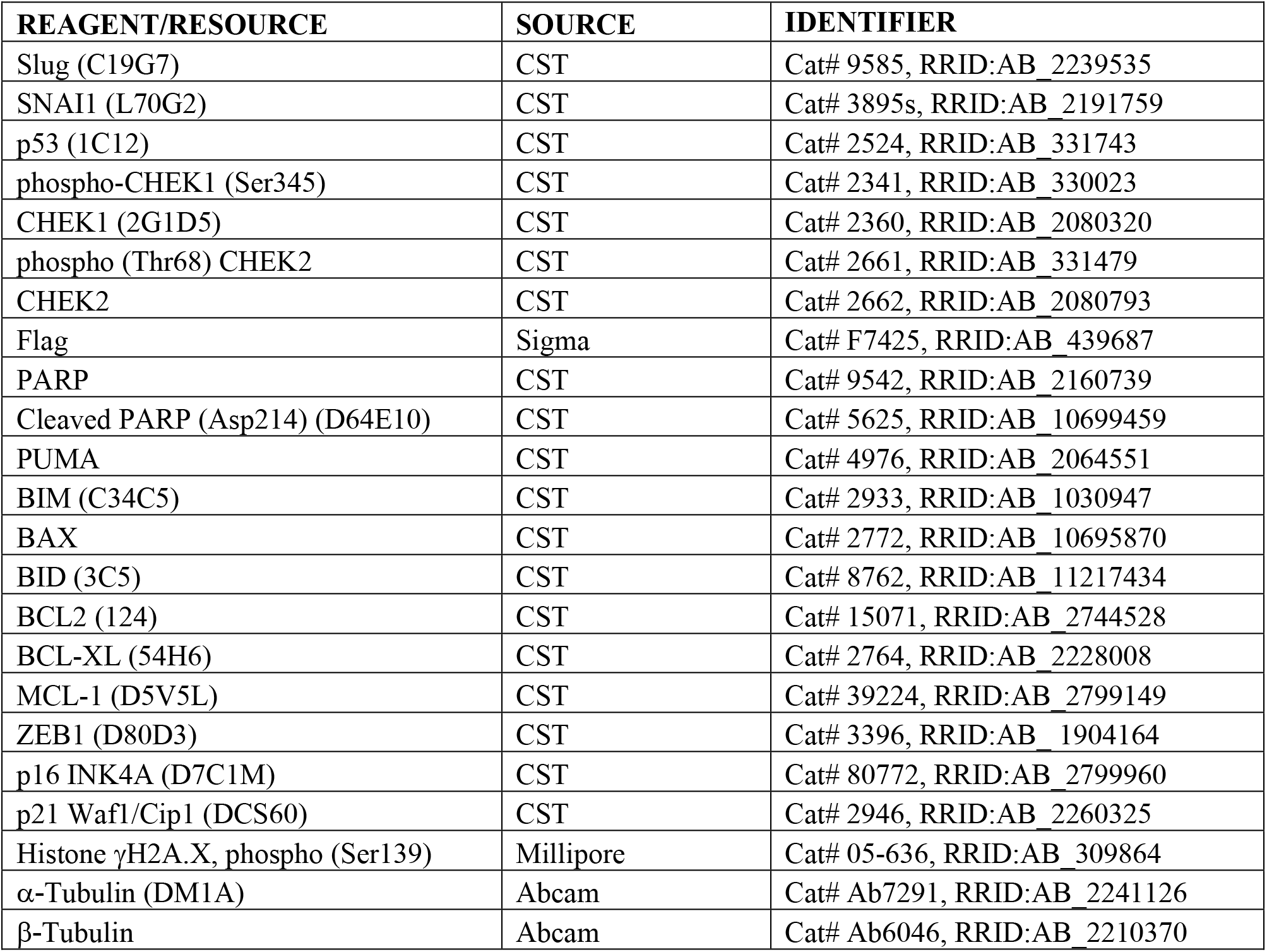
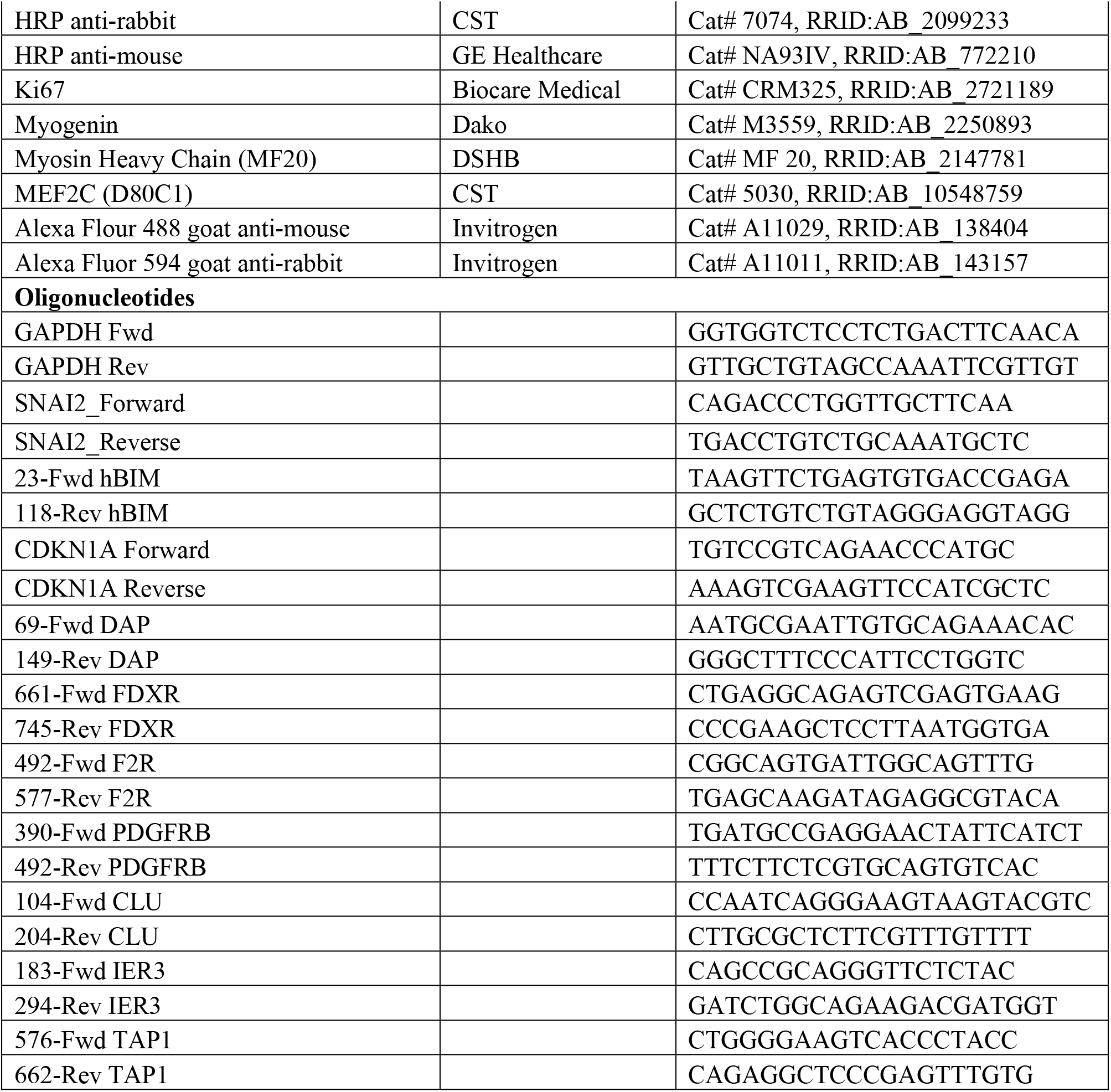

### SNAI2 protects RMS cells from IR *in vitro*

To assess whether *SNAI2* protects RMS cells from IR, we performed a *SNAI2*-knockdown experiment in Rh30, RD and Rh18 cells^24^, which were chosen as representative lines with low, moderate and high levels of radiosensitivity (Figure 1A-D). Compared to the scrambled control (Scr) shRNA, two *SNAI2* shRNAs (sh1 and sh2) reduced SNAI2 protein expression by 51-86% in Rh30, RD and Rh18 cells (Figures 2A, 2E, and 2I). Interestingly, while SNAI2 knockdown initially slowed cell proliferation^25^, this effect was no longer observed once stable lines were generated (Figure 2B, 2F, 2J). Each cell line was then exposed to an appropriate dose of IR (see Figure 1A-D) and analyzed for confluency every 4 hours for 5 days (Figure 2B, 2F, 2J). Compared to control-knockdown cells at 120 hours, *SNAI2*-knockdown cells became sensitized to IR across all three cell lines (Figure 2C, 2G, 2K; Rh30 Scr vs. sh1: *p*<0.0001, Rh30 Scr vs. sh2: *p*<0.0001; RD Scr vs. sh1: *p*<0.0001, RD Scr vs. sh2: *p*<0.0001; Rh18 Scr vs. sh1: *p*<0.0001, Rh18 Scr vs. sh2: *p*<0.0001; two-way ANOVA with Sidak’s multiple comparisons test). Similar results were observed in clonogenic colony-forming assays, where the surviving fraction of colonies was assessed in RMS cells exposed to a range of IR doses between 2 and 8 Gy. At lower doses of IR exposure, differences in survival and colony formation was minimal compared to higher doses of IR where there was a clear separation between the shScr and shSNAI2 treated cells (Figure 2D, 2H, 2L and Supplemental Figure 2A; 8 Gy Rh30 Scr vs. sh1: *p*<0.0001, 8 Gy Rh30 Scr vs. sh2: *p*<0.0001; 6 Gy RD Scr vs. sh1: *p*<0.0001, 6 Gy RD Scr vs. sh2: *p*<0.0001; 6 Gy Rh18 Scr vs. sh1: *p*<0.0001, 6 Gy Rh18 Scr vs. sh2: *p*< 0.0001; one-way ANOVA with Dunnett’s multiple comparisons test). Next, to test whether overexpression of SNAI2 could promote radioresistance, we transfected control pBabe vector and SNAI2-Flag constructs into the highly radiosensitive Rh18 cell line and tested the response to IR (Supplemental Figure 2B-D). While SNAI2 overexpression in Rh18 cells had no effect on proliferation in the absence of IR (Supplemental Figure 2C), the cells became less sensitive to 10 Gy IR at 120 hours (Supplemental Figure 2D; Rh18 pBabe vs. SNAI2-Flag: *p*<0.0001; two-way ANOVA with Sidak’s multiple comparisons test). Thus, SNAI2 protects RMS cells from IR *in vitro*.

**Figure 2.**
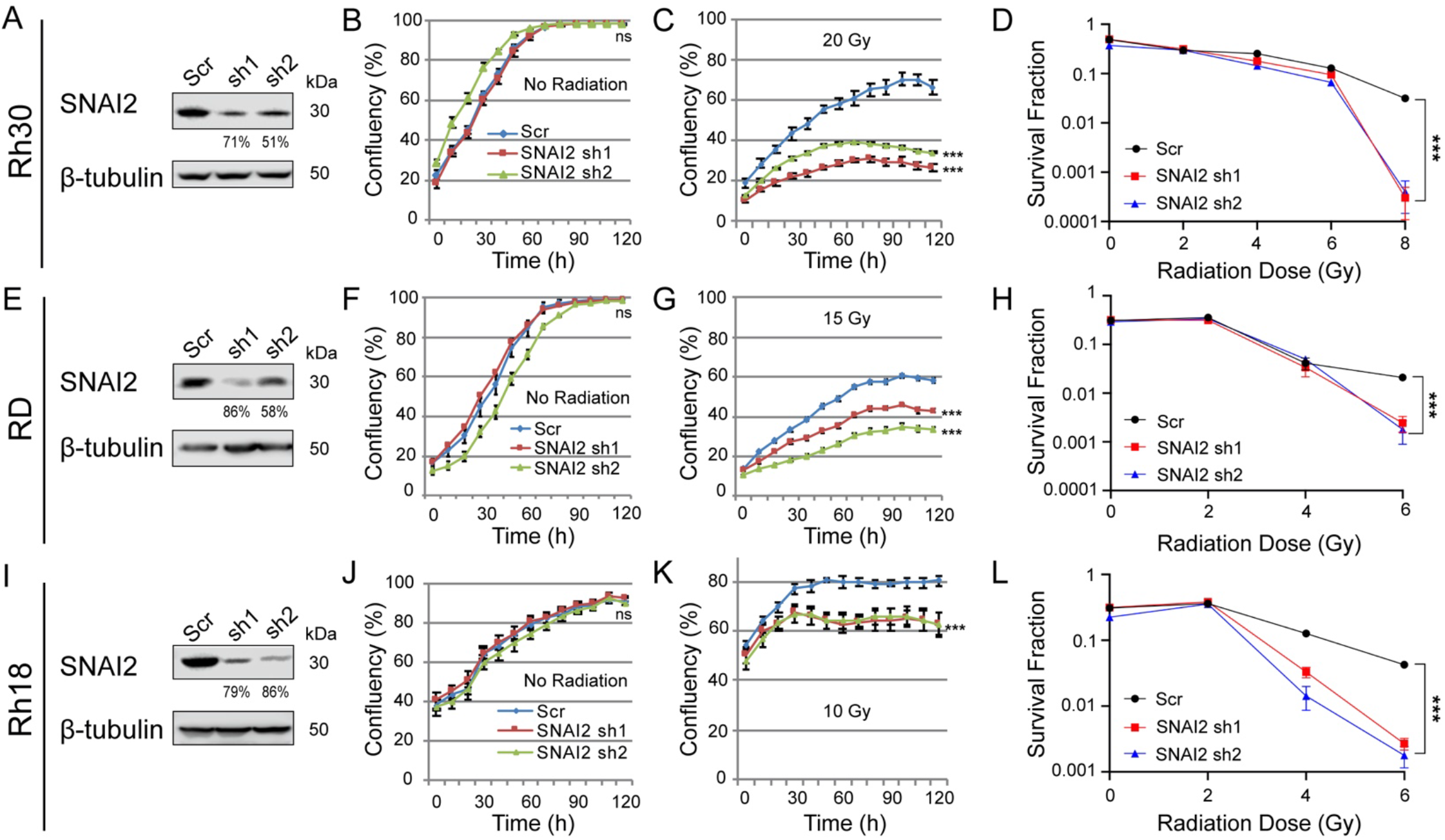
SNAI2 protects RMS cells from IR *in vitro.* A. Western blot showing protein expression for SNAI2 of control (Scr shRNA) or SNAI2 knockdown (sh1 or sh2) in Rh30 cells. B, C. Cell Confluency measured as a % of the total of Rh30 cells with no IR or IR at 20 Grays (Gy) with either control or SNAI2 knockdown was assessed using phase-contrast images acquired from 0h to 120 h. ns = not significant, \*\*\**p*<0.001. Error bars represent ±1 SD. D. Survival fractions of Rh30 Scr and SNAI2 knockdown colony formation assays were assessed at increasing IR dose exposures. Statistical differences were observed at 8 Gy. \*\*\**p*<0.0001. Error bars represent ±1 SD. E. Western blots showing protein expression for SNAI2 after control (Scr) or SNAI2 knockdown (sh1 or sh2) in RD cells. F. G. Cell Confluency measured as a % of the total of RD cells with no IR or IR at 15 Gy with either control or SNAI2 knockdown was assessed using phase-contrast images acquired from 0 to 120 h. ns = not significant, \*\*\**p*<0.001. Error bars represent ±1 SD. H. Survival fractions of RD Scr and SNAI2 knockdown colony formation assays were assessed at increasing IR dose exposures. Statistical differences were observed at 6 Gy. \*\*\**p*<0.0001. Error bars represent ±1 SD. I. Western blot showing protein expression of SNAI2 in control (Scr) or SNAI2 knockdown (sh1 or sh2) Rh18 cells. J. K. Cell Confluency measured as a % of the total of Rh18 cells with no IR or IR at 10 Gy in either control or SNAI2 knockdown cells was assessed using phase-contrast images acquired from 0 to 120 h. ns = not significant, \*\*\**p*<0.001. Error bars represent ±1 SD. L. Survival fractions of Rh18 Scr and SNAI2 knockdown colony formation assays were assessed at increasing IR dose exposures. Statistical differences were observed at 6 Gy. \*\*\**p*<0.0001. Error bars represent ±1 SD.

### SNAI2 protects RMS tumors from IR *in vivo*

We next questioned whether SNAI2 could protect RMS tumor cells from IR *in vivo.* We created murine xenografts of Rh30 and Rh18 cells with SNAI2 knockdown and Rh18 cells with SNAI2 overexpression, as well as appropriate controls, and performed irradiation experiments after each group of mice developed palpable tumors (200-400 mm^3^). Rh30 and Rh18 tumors with *SNAI2* (or control) knockdown were subjected to a cumulative 30-Gy dose of IR (2 Gy/day, 5 days per week, for 3 weeks). Following the completion of the IR regimen, tumors were analyzed weekly for relapse. Relapse was defined by re-growth of tumors to four times their size prior to IR treatment^26^. In the absence of IR, there were no significant differences in the growth rates between xenografts derived from Rh30 control- and *SNAI2*-knockdown cells (1×10^6^ cells injected/mouse, Figure 3A, Supplemental Figure 3A, n = 5). However, control Rh30 xenografts that were exposed to IR gave rise to relapse tumors significantly earlier (5 weeks post-IR) than *SNAI2*-knockdown Rh30 xenografts (11-14 weeks post-IR: Figure 3A, Supplemental Figure 3B, n = 8-10, *p*<0.001; Student’s t test). Similar to Rh30 xenografts, there were no differences in the growth rates between xenograft tumors derived from Rh18 control- and *SNAI2*-knockdown cells (5×10^6^ cells injected/mouse) in the absence of IR (Figure 3H, Supplemental Figure 3C), and post IR control Rh18 xenografts gave rise to relapsed tumors significantly earlier (7 weeks post-IR) than SNAI2-knockdown Rh18 xenografts (13-14 weeks post-IR: Figure 3H, Supplemental Figure 3D, n = 8-10, *p*<0.001; Student’s t test). Importantly, a subset of Rh18 SNAI2 shRNA knockdown tumors did not relapse as long as 21 weeks post IR. Finally, Rh18 xenografts (5×10^6^ cells injected/mouse) with control or SNAI2 overexpression showed no differences in growth rate in the absence of IR (Figure 3O, Supplemental Figure 3I, n = 4), but following a cumulative 20-Gy dose of IR (2 Gy/day, 5 days per week, for 2 weeks), SNAI2-overexpressing Rh18 xenografts relapsed more rapidly than control Rh18 xenografts (Figure 3O, Supplemental Figure 3J, n = 8-9, *p*<0.005; Student’s t test).

**Figure 3.**
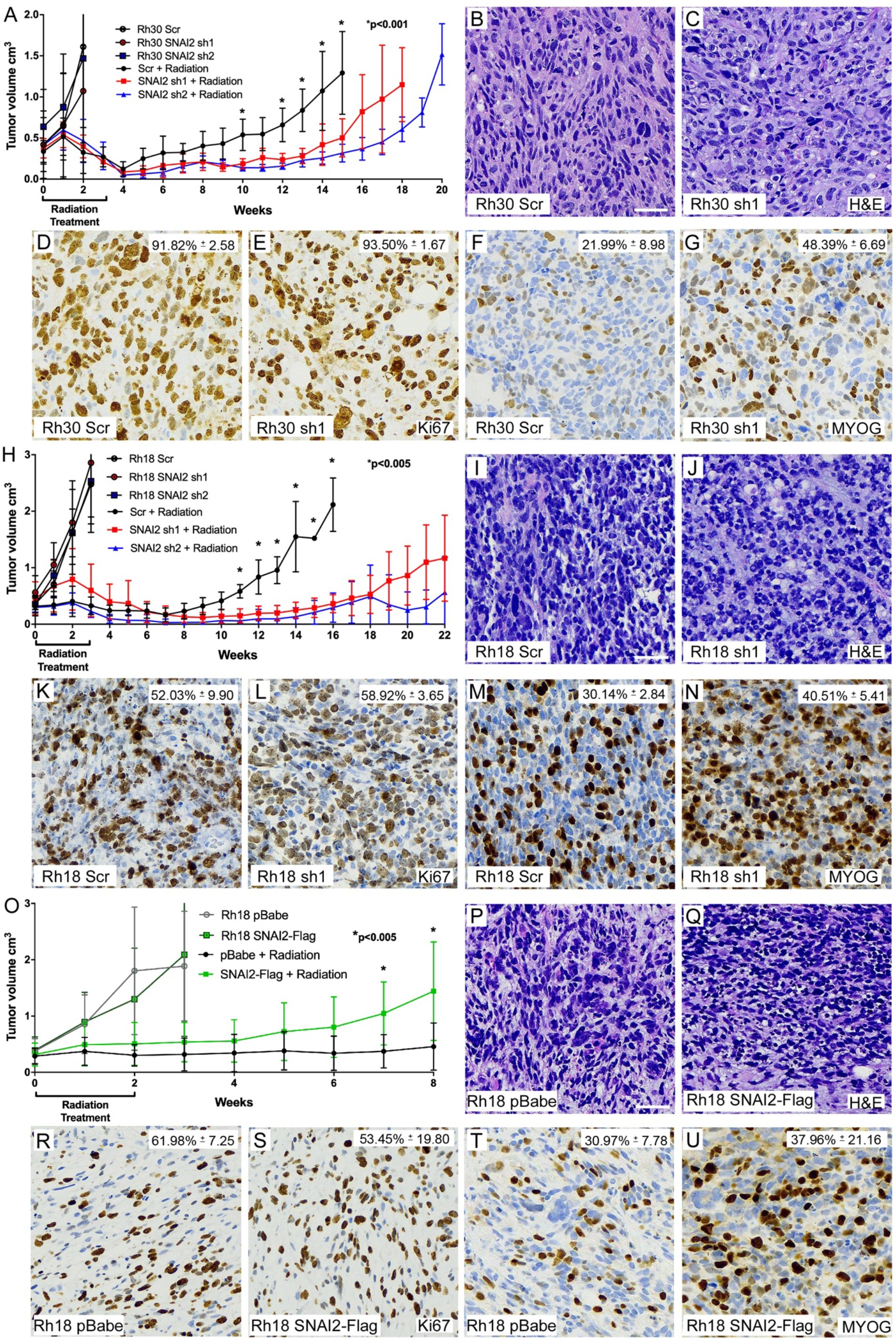
SNAI2 protects RMS tumors from IR *in vivo.* A. Growth curves of Rh30 xenografts including Scr shRNA and SNAI2 shRNA 1 and 2 (sh1, sh2) engrafted in mice. Xenograft growth was assessed under no IR and 30 Gy IR treatments. IR was given for 3 weeks at 2 Gy/day, 5 days a week. \**p*<0.001. Error bars represent ±1 SD. B, C. H&E staining showing histology of Rh30 Scr and SNAI2 knockdown tumor sections. Scale bar = 100 μm. D-G. Immunohistochemistry analysis of Ki67 (D, E) staining to assess proliferation and Myogenin (MYOG) staining (F, G) in Rh30 xenografts with either Scr shRNA or SNAI2 sh1. Ki67 –Scr vs. sh1 *p.* MYOG – Rh30 Scr vs. Rh30 sh1 *p*<0.05. Magnification same as B, C. H. Growth curves of Rh18 xenografts including Rh18 Scr shRNA and SNAI2 shRNA 1 and 2 (sh1, sh2) engrafted in mice. Xenograft growth was assessed under no IR and 30 Gy IR treatments. IR was given for 3 weeks at 2 Gy/day for 5 days a week. \**p*<0.005. Error bars represent ±1 SD. I, J. H&E staining showing histology of Rh18 Scr and SNAI2 knockdown tumor sections. Scale bar = 100 μm. K-N. Immunohistochemistry analysis of Ki67 (K, L) staining to assess proliferation and Myogenin (MYOG) staining (M, N) in Rh18 xenografts with either Scr shRNA or SNAI2 sh1. MYOG – Scr vs. sh1 *p*=0.0519. Magnification same as I, J. O. Growth curves of Rh18 xenografts expressing control vector (pBabe) and SNAI2-Flag engrafted in mice. Xenograft growth was assessed under no IR and 20 Gy IR treatments. IR was given for 2 weeks at 2 Gy/day for 5 days a week. \**p*<0.005. Error bars represent ±1 SD. P. Q. H&E staining showing histology of Rh18 pBabe and SNAI2-Flag tumor sections. Scale bar = 100 μm. R-U. Immunohistochemistry analysis of Ki67 (R, S) staining to assess proliferation and Myogenin (MYOG) staining (T, U) in Rh18 xenografts with either pBabe or SNAI2-Flag expression. Ki67 – pBabe vs. SNAI2-Flag not significantly different. MYOG – pBabe vs. SNAI2-Flag not significantly different. Magnification same as P,Q.

To investigate the effect of *SNAI2* knockdown on tumor histology, proliferation and differentiation in Rh30 and Rh18 relapsed tumors, xenografts with Scrambled or *SNAI2* knockdown were harvested from mice once they reached 4X their initial volume and tumors were processed, sectioned and assessed for histology (H&E), proliferation (Ki67), and differentiation (MYOG). H&E analysis did not show significant differences between Scrambled vs. SNAI2-knockdown tumors (Figure 3B-C, 3I-J). Next, Ki67 staining showed Control (shScr and SNAI2 knockdown xenografts proliferate at similar rates (Figure 3D-E, 3K-L). In Rh30 and Rh18 tumors with SNAI2 shRNA knockdown, there was a trend toward areas showing increased MYOG expression (Figure 3F-G, 3M-N; Rh30 Scr vs. Rh30 sh1 *p*<0.05, Rh30 Scr vs. Rh30 sh2 *p*<0.05; Rh18 Scr vs. Rh18 sh1 *p*=0.0519, Rh18 Scr vs. sh2 *p*=0.0852). However, this increase in MYOG did not result in terminal differentiation as assessed by MyHC (MF20) staining (Supplemental Figure 3E-H). Similarly, there was not a significant effect on tumor histology, proliferation, or differentiation observed when comparing Rh18 controls to Rh18 SNAI2 overexpressing xenografts (Figure 3P-U; Rh18 pBabe vs. SNAI2-Flag not significant; Welch’s t-test). Thus, while the H&E and MYOG staining showed that SNAI2 may also inhibit myogenic differentiation in some RMS tumors; however, this effect is not as prominent compared to our previously described results in RAS mutant ERMS^25^. Altogether, our data suggest that the major conserved effect of SNAI2 post IR is to protect RMS tumors from IR *in vivo*.

### Loss of SNAI2 promotes IR-mediated apoptosis and blocks irradiated RMS cells from exiting the cell cycle

To better understand how SNAI2 protects RMS cells from IR, we assessed the effects of SNAI2 knockdown on apoptosis and the cell cycle in RMS cell lines. We first analyzed live cells for signs of apoptosis using a Caspase-Glo assay in Rh30, RD and Rh18 cells. In the absence of IR, control- and SNAI2-knockdown cells grew at similar rates (see Figure 2B, 2F, 2J) and had significantly lower Caspase3/7 staining until they reached confluence in all three RMS cell lines (Supplemental Figure 4A). However, at 72 hours post-IR (hpIR), SNAI2-knockdown cells exhibited a significant increase in apoptosis compared to controls (Figure 4A-C, Supplemental Figure 4B; 15 Gy Rh30 Scr vs. sh1 *p*<0.0001; 10 Gy RD Scr vs. sh1 *p*=0.001; 5 Gy Rh18 Scr vs. sh1 *p*< 0.0001; two-way ANOVA with Sidak’s multiple comparison). We then performed independent apoptosis assays using annexin V/propidium iodide staining to confirm these findings by analyzing live cells post IR by flow cytometry. Consistent with the Caspase-Glo assay, there was a significant increase in early and late apoptotic cells in the SNAI2-knockdown Rh30, RD and Rh18 populations between 72 to 120 hpIR compared to control-knockdown cells (Figure 4D-G, Supplemental Figure 4C-D; Rh30 15 Gy/120 hpIR, early apoptosis, *p*<0.0001, late apoptosis, *p*<0.0001; RD 10 Gy/72hpIR, early apoptosis, *p*<0.0001, late apoptosis, *p*<0.0001; Rh18 5 Gy/96 hpIR, early apoptosis, *p*<0.0001, late apoptosis, *p*<0.0001; Two Proportions Z-test). Importantly, in the absence of IR, there were no significant differences in apoptosis between control- and SNAI2-knockdown cells for each of the cell lines (Supplemental Figure 4G-H, 4K-L, 4O-P). These experiments indicate that SNAI2 protects RMS cells from IR-induced apoptosis.

**Figure 4.**
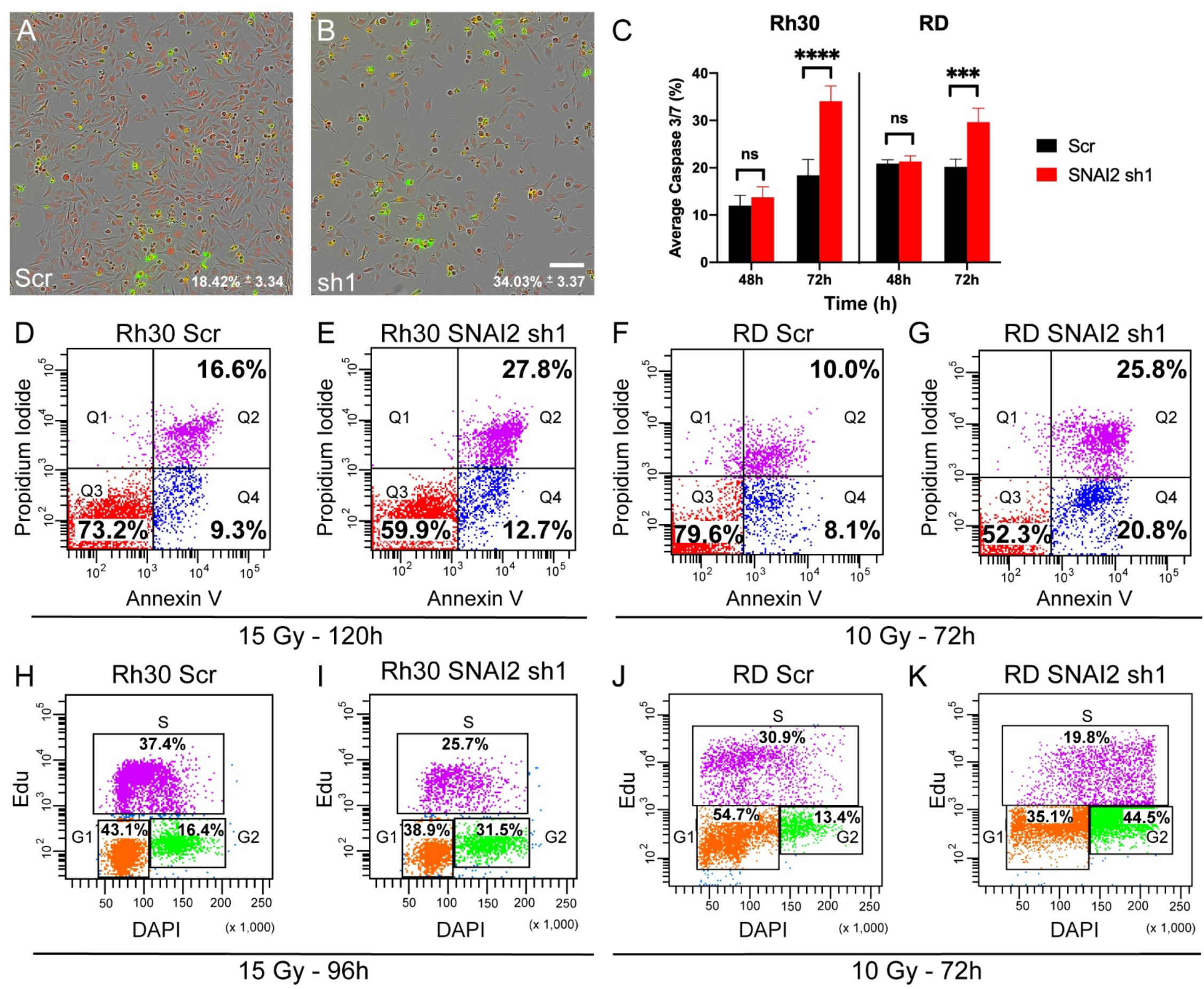
Loss of SNAI2 promotes IR-mediated apoptosis and blocks irradiated RMS cells from exiting the cell cycle. A, B. Representative images of Caspase-Glo assay in Rh30 cells (either control or SNAI2 knockdown) at 72h post IR exposure (15 Gy) with red labeling cells/nuclei and green labeling caspase 3/7; average caspase 3/7 levels (%) were quantified in C. Scale bar = 150 μm. C. Average Caspase 3/7 percentage (mean ± 1 SD) in Rh30 and RD Scr and SNAI2 sh1 cells 48h and 72h after IR exposure of 15 or 10 Gy respectively. \*\*\**p*<0.001, \*\*\*\**p*<0.0001. D-G. Flowcytometry plots showing Propidium iodide vs. Annexin V staining of Rh30 and RD Scr or SNAI2 sh1 cells and treated with indicated IR doses. Q4 represents cells undergoing early apoptosis, whereas Q2 represents cells undergoing late apoptosis. Q3 represents live cells not undergoing apoptosis. Rh30 early apoptosis (Q4): Scr 9.3% vs. 12.7%, *p*<0.0001, late apoptosis (Q2): Scr 16.6% vs. sh1 27.8%, *p*<0.0001; RD early apoptosis (Q4): Scr 8.1% vs. sh1 20.8%, p<0.0001, late apoptosis (Q2): Scr 10.0% vs. sh1 25.8%, *p*<0.0001. H-K. Flowcytometry plots of EdU vs. DAPI staining in Rh30 and RD cells with either Scr shRNA or SNAI2 sh1 after exposure to indicated IR doses. Rh30 Scr vs. sh1 G2 phase *p*<0.0001; RD Scr vs. sh1 G2 phase *p*<0.0001.

We next analyzed whether SNAI2 regulates the cell cycle after exposure to IR using EdU labeling followed by flow cytometry. In the absence of IR, there were no differences in proliferation between control- and SNAI2-knockdown cells (Supplemental Figure 4I-J, 4M-N, 4Q-R; Rh30 Scr vs. sh1, *p*=0.7062; RD Scr vs. sh1, *p*=1; Rh18 Scr vs. sh1, *p* = 0.9228; Two Proportions Z-test). However, following IR treatment, SNAI2-knockdown cells showed a significant reduction in cells in the G1 and S phases and an accumulation of cells in the G2-M phase of the cell cycle, indicative of a G2/M block (Figure 4H-K, Supplemental Figure 4E-F; Rh30 15 Gy/96h, Scr vs. sh1, *p*<0.0001; RD 10 Gy/72h, Scr vs. sh1, *p*<0.0001; Rh18 5 Gy/96h, vs. sh1, *p*<0.0001; Two Proportions Z-test). These experiments suggest that loss of SNAI2 may prevent mitosis or alter progression through the M-phase of cell cycle following exposure to IR.

### SNAI2 represses the expression of the BH3-only gene *BIM* in RMS cells

To investigate the mechanisms by which *SNAI2* protects RMS cells from IR-induced apoptosis, we analyzed the expression of both pro- and anti-apoptotic regulators of mitochondrial apoptosis in Rh30, RD, and Rh18 cells (Figure 5A-B, Supplemental Figure 5A). We confirmed that in all three cell lines, SNAI2 knockdown persisted for the duration of the experiment. In response to IR, SNAI2-knockdown cells showed increased expression of the apoptosis marker cleaved-PARP1, which is prominent especially between 72 to 96 hpIR and consistent with the annexin-V and Caspase-Glo analyses (Figure 4D-G, Supplemental Figure 4C-D). In the absence of IR, expression of the proapoptotic BH3-only gene *PUMA/BBC3* appears to be upregulated upon SNAI2 knockdown in all cell lines. Interestingly, SNAI2 knockdown also elicited a prominent increase in the proapoptotic BH3-only BIM across all three cell lines as well as in BID and BAX in Rh18 and Rh30 lines. With respect to anti-apoptotic regulators, BCL-2 showed a modest increase in both Rh30 and Rh18 in response to SNAI2 knockdown in the absence of IR. MCL-1 expression is slightly decreased only in Rh30, and BCL-xl expression was elevated in RD and Rh30. Moreover, in response to IR treatment, BCL-2, MCL-1, and BCL-xl varied across cell lines and failed to correlate with SNAI2 knockdown. Among the regulators of apoptosis, only BIM expression was found to be consistently elevated across all cell lines in response to SNAI2 knockdown in IR-exposed cells. These experiments indicate that SNAI2, in addition to its known function as a transcriptional repressor of PUMA^10, 27^, appears to repress the expression of BIM in RMS cells.

**Figure 5.**
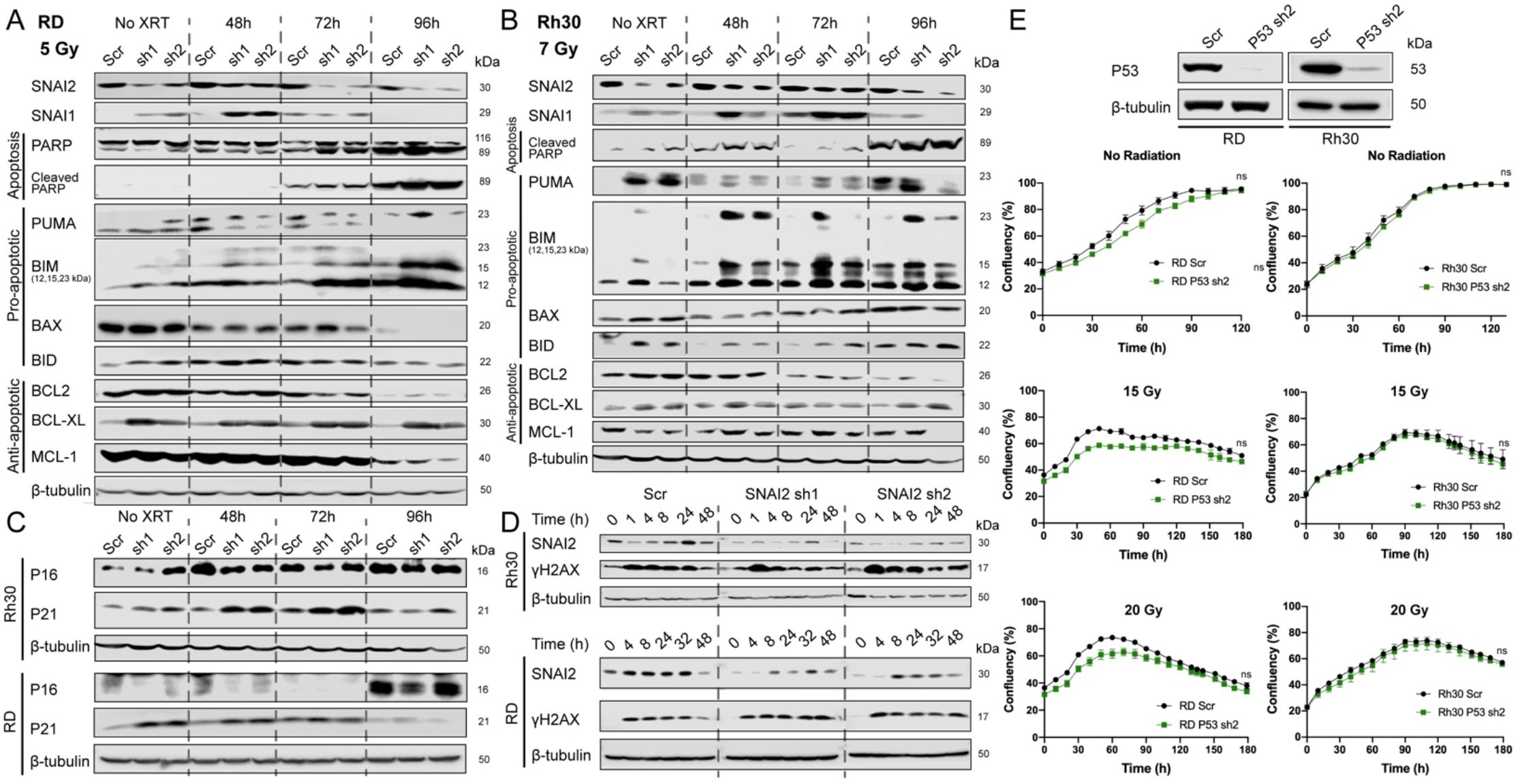
SNAI2 represses the expression of the BH3-only gene BIM in RMS cells. A, B. Western blot analyses to determine protein expression of SNAI2, SNAI1, PARP, Cleaved PARP, PUMA, BIM, BAX, BID, BCL2, BCL-XL, and MCL-1 in RD and Rh30 cells under no IR or at 48-, 72-, or 96-hours post-IR at 5 Gy (RD) and 7 Gy (Rh30) treatments. C. Western blot analysis to determine protein expression of P16 and P21 in Rh30 and RD cells under no IR or at 48-, 72-, or 96-hours post-IR with 7 Gy (Rh30) and 5 Gy (RD). D. Western blot analysis of gH2AX over time after exposure to IR in Rh30 (7 Gy) and RD (5 Gy) control (Scr) and SNAI2 knockdown (sh1 and sh2) cells. E. Western blot analysis to determine protein levels of P53 in RD and Rh30 cells (either Scr control or P53 sh2 knockdown). Confluency (%) of non-IR or IR-treated (15 or 20 Gy) RD and Rh30 cells (with either Scr control or P53 shRNA knockdown) was assessed on phase-contrast images acquired from 0 to 180 h. No statistical differences were observed. ns = not significant. Error bars represent ±1 SD.

To determine the mechanism by which SNAI2 influences the cell cycle in RMS cells, we analyzed the expression of the *P21/CDKN1A* cell cycle checkpoint inhibitor and found that SNAI2 knockdown leads to upregulation of P21 in both Rh30 and RD cells (Figure 5C). Since P21 is also a marker of cells undergoing senescence, we analyzed P16 expression and performed β-gal staining in Rh30 and RD cells. There were no differences in P16 expression or β-gal staining between control and SNAI2 knockdown in either cell type, suggesting that senescence is not regulated by SNAI2 in these cells (Figure 5C, Supplemental Figure 5B-E). In RMS tumors, *CDKN1A/P21* expression is often repressed, and re-expression of P21 promotes differentiation ^28, 29^. Moreover, SNAI2 has been shown by ChIP-seq experiments to indirectly block *CDKN1A* expression in ERMS RD and SMS-CTR cells, and co-knockdown of *CDKN1A* and *SNAI2* in RD and JR1 RMS cells results in the loss of differentiation-positive, myosin-heavy-chain-expressing cells^30^. We also assayed the effect of SNAI2 knockdown on differentiation post IR in Rh30 and RD cells and in another ERMS cell line with wild-type P53, Rh36. Both RD and Rh36 are RAS mutant ERMS cell lines. Following exposure to IR, RD and Rh36 cells with SNAI2 knock down exhibited a significant increase in differentiation as determined by the expression of differentiated myosin MF-20 staining (Supplemental Figure 5F-K; MF-20: RD Scr vs. sh1 *p*<0.0001, RD Scr vs. sh2 *p*<0.005, Rh36 Scr vs. sh1 *p*<0.0001; MEF2C: RD Scr vs. sh1 *p*<0.0001, RD Scr vs. sh2 *p*<0.005, Rh36 Scr vs. sh1 *p*<0.0001; two-way ANOVA with Sidak’s multiple comparisons test). In contrast, in Rh30 (ARMS) cells this effect was not as prominent (Supplemental Figure 5L, MF-20: Rh30 Scr vs. sh1, not significant, Rh30 Scr vs. sh2, not significant; MEF2C: Rh30 Scr vs. sh1 *p*<0.005, Rh30 Scr vs. sh2, *p*<0.0001; two-way ANOVA with Sidak’s multiple comparisons test), suggesting that SNAI2 may be more important for the suppression of muscle differentiation in RAS mutant ERMS tumors consistent with our previous findings^25^.

We next questioned whether SNAI2 regulates IR-induced DNA repair. Since IR primarily causes DNA double-stranded breaks, we analyzed the expression of a well-established marker for these DNA lesions, yH2AX^31^, during a time course following IR in Rh30 and RD cells via western blot analysis (Figure 5D). As expected, γH2AX levels increased rapidly following exposure to IR in both cell lines with SNAI2-knockdown cells showing similar increases compared to control-knockdown cells; however, γH2AX expression was retained as late as 48h in SNAI2 knockdown cells, suggesting a delay in the ability of SNAI2 knockdown cells to repair damaged DNA (Figure 5D). However, SNAI2-knockdown RD and Rh30 cells have little if any differences in expression of DNA damage checkpoint regulators pCHEK1 and pCHEK2 following IR (Supplemental Figure 5M-N). These findings suggest that SNAI2 might have additional roles on influencing the timing of repair of DNA double-strand breaks in RMS cells.

Both Rh30 and RD cells have *TP53* mutations that aberrantly stabilize the P53 protein (Figure 1E)^32, 33^. To determine if the expression of mutant P53 is important for IR-mediated inhibition of cell growth in RD and Rh30 cells, we performed an shRNA-mediated knockdown of P53 and tested its effects on the RD and Rh30 lines (Figure 5E). In both lines, knockdown of mutant P53 failed to affect sensitivity to IR (Figure 5E). Additionally, in both Rh30 and RD cells, SNAI2 is slightly induced or maintained following IR suggesting alternate mechanisms by which SNAI2 expression is modulated in RMS (Figure 5D).

### Direct repression of *BIM* by SNAI2 blocks apoptosis in irradiated RMS cells

Since BIM was consistently and robustly induced by loss of SNAI2 in RMS cells and apoptosis post IR is the major effect of SNAI2 ablation across all RMS cells assessed, (Figure 5A-B, Supplemental Figure 5A) we questioned whether SNAI2 could directly repress expression of the *BIM.* We first performed RNA-seq analysis comparing control- and SNAI2-knockdown in RD cells at 24 hpIR (5 Gy). GSEA pathway analysis revealed that in addition to differences in myogenic differentiation, several stress response pathways were modulated in response to IR. These pathways included “hypoxia”, “UV response down”, and “apoptosis” (Figure 6A). Several apoptotic pathway genes were increased or decreased in SNAI2 knockdown cells post IR (Figure 6B). We performed qRT-PCR analysis using RD and Rh30 cells for a subset of the apoptotic regulators and validated the upregulation of *CDKN1A* (only RD), *BCL2L11(BIM),* and *DAP* and downregulation of *FDXR, F2R, PDGFRB, CLU, IER3,* and *TAP1* (Figure 6C). Next, ChIP-seq analysis of SNAI2 binding in RD and SMS-CTR cells using both control- and SNAI2-knockdown treatments revealed SNAI2-binding peaks in the enhancer regions of *DAP* (Supplemental Figure 6A) and *BCL2L11/BIM* (Figure 6D)^25^. However, no peaks were found associated with the promoter enhancer regions of *CDKN1A* (Supplemental Figure 6B)^25^. Moreover, we observe that the SNAI2 peaks in *BIM/BCL2L11* were lost upon *SNAI2* knockdown and is associated with increased expression of *BIM/BCL2L11.* These experiments establish that SNAI2 is likely a direct repressor of pro-apoptotic *BIM*/*BCL2L11*.

**Figure 6.**
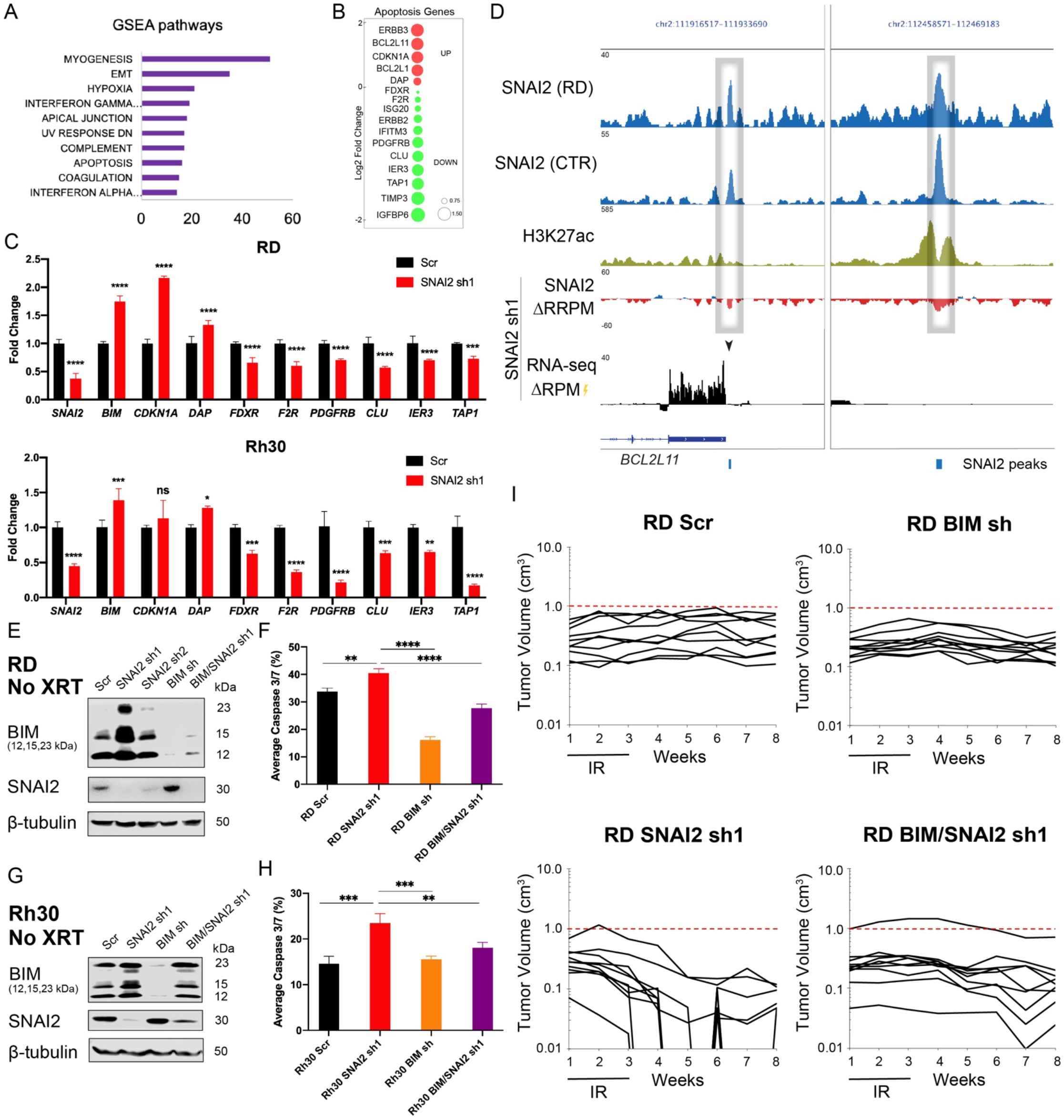
Direct repression of *BIM* by SNAI2 blocks apoptosis in irradiated RMS cells. A. GSEA pathway analysis comparing mRNA expression (RNA-seq) in control/Scr and SNAI2 shRNA treated RD cells 24 h post IR (5 Gy). Enriched pathways in shSNAI2 cells (GSEA Hallmark pathways) with number of up-regulated genes in each pathway class (x-axis) B. RNA-seq data showing genes highly upregulated or downregulated when comparing control/ Scr- and SNAI2-shRNA knockdown RD cells 24 h post IR (5 Gy). Red circles denote up-regulated genes and Green denotes down-regulated genes; size of circles represent Log2 fold change compared to shScr. C. Real time qPCR analysis (mean ± 1 SD) of various cell cycle and apoptosis genes in RD and Rh30 cells 48h after irradiation with 5 Gy and 10 Gy, respectively. ns = not significant, \**p*<0.05, \*\**p*<0.005, \*\*\**p*<0.001, \*\*\*\**p*<0.0001. D. ChIP-seq tracks of SNAI2 and H3K27ac binding at the *BIM/BCL2L11* locus in RD and SMS-CTR cells in control /Scr, delta (Δ) enrichment value (shSNAI2 sh1 minus shScr) for SNAI2 and gene expression (RNA-seq). Boxed area corresponds to SNAI2 binding region and blue line represents SNAI2 called peak. RRPM, Reference-adjusted Reads Per Million Mapped Reads; RPM, Reads Per Million Mapped Reads. E. Western blot analyses of BIM and SNAI2 expression in non-IR treated RD cells under either Scr control, SNAI2 shRNA, BIM shRNA, or double BIM/SNAI2 shRNA conditions. F. Average Caspase 3/7 (%) (mean ± 1 SD) in RD Scr, SNAI2 sh1, BIM sh, and BIM/SNAI2 sh1 cells 72h after IR exposure (15 Gy). \*\**p*<0.005, \*\*\*\**p*<0.0001. G. Western blot analyses of BIM and SNAI2 expression in non-IR treated Rh30 cells under either Scr control, SNAI2 shRNA, BIM shRNA, or double BIM/SNAI2 shRNA conditions. H. Average Caspase 3/7 (%) (mean ± 1 SD) in Rh30 Scr, SNAI2 sh1, BIM sh, and BIM/SNAI2 sh1 cells 72h after IR exposure (15 Gy). \*\**p*<0.005, \*\*\**p*<0.001. I. Individual tumor volumes of RD Scr, SNAI2 sh1, BIM sh, and BIM/SNAI2 sh1 xenografts after 30 Gy (2 Gy/day, 5x a week, for 3 weeks). Red dashed line indicates 1 cm^3^.

We next questioned whether reducing BIM expression would abrogate the SNAI2-knockdown-induced radiosensitivity in RMS cells. We therefore generated RD and Rh30 cells with stable lentiviral infections for 1) control shRNA, 2) *SNAI2* shRNA, 3) *BIM* shRNA and 4) both *SNAI2* and *BIM* shRNAs^34^. Consistent with our earlier findings, knockdown of SNAI2 increased the levels of all three forms of BIM, which was reversed by concomitant knockdown of BIM (Figure 6E, 6G). As expected, IR treatment of SNAI2-knockdown RD and Rh30 cells with a single dose of 15 Gy resulted in selective loss of confluency (Supplemental Figure 6C-D) and a significant increase in Caspase 3/7 positive cells compared to irradiated control-knockdown cells. However, this effect on apoptosis and confluency was significantly reversed in SNAI2/BIM-double knockdown cells post-IR (Figure 6F, 6H, Supplemental Figure 6C-F; Caspase 3/7: RD SNAI2 sh1 vs RD BIM/SNAI2 sh1 *p*<0.0001; Rh30 SNAI2 sh1 vs Rh30 BIM/SNAI2 sh1 *p*=0.0098; one-way ANOVA with Tukey’s multiple comparisons test. Confluency at 110 hp IR: RD BIM/SNAI2 vs. SNAI2 sh1 *p*<0.0001; Rh30 BIM/SNAI2 vs. SNAI2 *p*<0.0001; two-way ANOVA with Sidak’s multiple comparisons test). Of note, in the absence of IR, no differential effects on confluency or apoptosis was observed across all four groups (Supplemental Figure 6D-F). Since the SNAI2/BIM-knockdown-mediated rescue of radiosensitivity in Rh30 cells was only partial, it is possible that other SNAI2-regulated genes may also contribute to IR-induced effects (Supplemental Figure 6D). Finally, we tested in RD xenografts the effect of combined knockdown of SNAI2 and BIM on response to IR. We created murine xenografts of RD with shScr, shBIM, shSNAI2, and shSNAI2/BIM double knockdown conditions. After each group of mice developed palpable tumors (200-400 mm^3^), they were subjected to a cumulative 30-Gy dose of IR (2 Gy/day, 5 days per week, for 3 weeks). Following the completion of the IR regimen, tumors were analyzed weekly for changes in tumor volume. While shScr and shBIM tumors showed no effect on relative change in tumor volume post-IR compared to their initial volumes (Change in volume 2 weeks post-IR: shScr = 0.0295 cm^3^/week, shBIM = 0.01125 cm^3^/week; no significant difference, One-way ANOVA with Tukey’s multiple comparisons test), in the shSNAI2 group (Figure 6I top panels), 7 of 10 tumors showed a complete response, while the other 3 of 10 show a partial response 4 weeks post-IR. In contrast, in the double SNAI2/BIM shRNA group only 2 of 8 tumors resulted in a complete response, whereas 6 of 8 tumors had only a partial response to IR treatment. Indeed, the relative change in tumor volume post-IR compared to their initial volumes in shSNAI2 xenografts were significantly decreased compared to double SNAI2/BIM knockdown xenografts (Figure 6I bottom panels) (Change in volume 2 weeks post-IR: shSNAI2 = −0.067 cm^3^/week, shSNAI2/BIM = – 0.0105 cm^3^/week; *p*<0.0001, One-way ANOVA with Tukey’s multiple comparisons test). Together, these experiments show that SNAI2-mediated repression of *BIM* protects RMS cells from the effects of IR.

## DISCUSSION

Radiation therapy is an important component of RMS treatment, especially during the management of metastases in high-risk patients^2, 3, 5^. Identifying pathways that regulate the response to IR therapy could potentially provide biomarkers of resistance or sensitivity to IR as well as targets for therapeutic intervention. Here, we show that SNAI2 directly represses the proapoptotic BH3-only gene *BIM* to protect RMS tumors from IR.

SNAI2 protects RMS tumor cells from IR despite the fact that the P53 pathway is nonfunctional or is mutant and consequently PUMA induction is not robust in the *TP53* mutant RMS cell lines tested in this study. In lymphocytes and other highly radiosensitive cell types, P53 becomes rapidly activated in response to IR, and triggers mitochondrial apoptosis through induction of the expression of *PUMA*^10^. Our finding that SNAI2 expression levels appear to dictate radiosensitivity of RMS cells in a manner that is independent of *TP53* mutant status and possibly PUMA expression, but rather by repression of BIM that is not directly regulated by P53, suggests an important role for SNAI2 in the radiation response in cancer cells that can be dependent or independent of P53. Experiments *in vivo* in mice mutant for *Puma* and *Bim* and in cell lines and malignant cells demonstrate that both Puma and Bim can potently trigger the mitochondrial apoptosis response post radiation and that loss of Bim protects lymphocytes from radiation and also decreases the time to tumor initiation in thymocytes compared to wild-type controls^11, 35–37^ This effect of BIM protecting cells from radiation is seen not only in renal cell carcinomas and also observed by correlation in a study in KRAS mutant lung cancer cell lines. For example, renal cell carcinomas often have mutations in VHL and these tumors express low levels of BIM (EL) and are more resistant to several apoptotic stimuli, including UV-radiation^38^. Also, in a subset of KRAS mutant lung cancer cell lines, low BIM expression was associated with relative resistance to radiation^39^. Thus, BIM is a bona fide BH3 pro-apoptotic regulator that can be induced post IR to mediate mitochondrial apoptosis. Our study shows that in RMS tumors SNAI2 is a potent repressor of *BIM* expression, yet its expression can be independent of *TP53.* In support of this assertion, we have recently found in the ERMS sub-type that MEK signaling and MYOD maintains SNAI2 expression; while in the ARMS subtype the PAX3-FOXO fusion oncogene is also known be required for *SNAI2 e*xpression^40, 41^.

Despite the presence of different genetic drivers namely, PAX3-FOXO fusion oncogene in ARMS lines and mutant RAS and amplified MDM2 in the ERMS lines among the cell lines used in this study, the radiosensitivity of each cell line correlated with expression of SNAI2 rather than oncogene/tumor suppressor status. This suggests that *SNAI2* expression levels could be used to predict the degree of radiosensitivity across different RMS tumors, and perhaps multiple tumor types. For example, most solid tumors have intermediate levels of *SNAI2* and intermediate sensitivities to IR^42, 43^ (Supplemental Figure 1C). In contrast, melanomas and osteosarcomas, which are generally not treated with IR in the clinic due to their inherent radioresistance and radiation when administered is at high doses for local control of disease and in palliative care^44–46^, have the highest levels of *SNAI2* expression. This makes sense from a developmental perspective since sarcomas originate from mesoderm^47, 48^, a tissue that expresses high levels of SNAI2 during development and has roles in muscle, bone and cartilage tissues (Reviewed in^8^). The analysis of SNAI2 expression levels could therefore be potentially informative for the treatment of multiple different cancers.

RMS cells with stable SNAI2 knockdown while showing relatively little differences in proliferation, apoptosis or cell viability compared to controls, nevertheless have consistent increased expression of pro-apoptotic modulators BIM, PUMA and BID at baseline even in cells not exposed to IR. Additionally, our studies show that SNAI2 directly represses *BIM* expression. This suggests that SNAI2 knockdown cells are primed for mitochondrial apoptosis and an important role oncogenic role for SNAI2 is to prevent IR induced mitochondrial apoptosis. In contrast to the consistent expression of BIM and PUMA in untreated cells different anti-apoptotic regulators, BCL2, BCLxl and MCL-1, are expressed at varying levels in the Rh30, RD, and Rh18 RMS lines to balance pro apoptotic factor expression in SNAI2 knockdown cells. Based on these observations, one would predict that treating RMS cells with inhibitors of BCL2, BCLxl, and MCL1 in the context of SNAI2 knockdown would have variable effects on mitochondrial apoptosis and might be an important factor when considering combination treatments.

In summary, our study implicates SNAI2 as a potential biomarker for IR sensitivity in RMS, with an inverse correlation between SNAI2 expression levels and radiosensitivity of the tumor cells. Post IR in conditions where SNAI2 expression is reduced, both ERMS and ARMS cell lines exhibit significantly reduced cell growth in addition to increased levels of apoptosis. ERMS cell lines may also undergo differentiation following exposure to IR. Differences in other known pathways that could explain reduced cell proliferation, such as senescence and dysfunctional DNA repair, were either variable or not observed in the SNAI2-knockdown cells compared to control RMS cells. Indeed, the finding that SNAI2 directly represses BIM, a potent inducer of mitochondrial apoptosis, supports the existence of an exploitable SNAI2/BIM signaling axis in RMS or potentially other tumors with high SNAI2 expression, which could ultimately improve the efficacy of IR therapy in the clinic.

## Materials and Methods

### Animals

Animal studies were approved by the University of Texas – Health San Antonio Committee on Research Animal Care under protocol #20150015AR. C.B-Igh-1b/IcrTac-Prkdcscid (SCID) female mice, aged 6-8 weeks, were used for *in vivo* xenograft experiments.

### Mouse xenograft and *in vivo* IR experiments

Rh30, Rh18, and RD cells with scrambled (shScr), SNAI2 knockdown (shSNAI2) and RD cells with scrambled (shScr), SNAI2 knockdown (shSNAI2), BIM knockdown (shBIM)- and double BIM/SNAI2 knockdown (shBIM/SNAI2)- treatment conditions were collected, counted, and analyzed by flow cytometry to determine viability using DAPI. Equal numbers of viable cells were then embedded into Matrigel at a final concentration of 1×10^6^ (Rh30) or 5×10^6^ (Rh18) cells per 100 μl and injected subcutaneously into anesthetized mice. Tumor growth was monitored and measured weekly using a caliper scale to measure the greatest diameter and length, which were then used to calculate tumor volume. While a subset of tumors was monitored without any treatment, another subset was subjected to low-dose IR therapy for 3 weeks (2 Gy/day; 5 days a week), receiving a total of 30-Gy IR (PXi Precision X-Ray X-RAD 320). Tumor volume was monitored throughout the treatment and during the weeks following treatment. Comparisons between groups was performed using a Student’s t test. Rh18 pBabe and SNAI2-Flag xenografts were performed as above, with the treatment arm receiving 2 weeks of IR therapy (2 Gy/day for 5 days/week) for a total of 20 Gy.

### Human rhabdomyosarcoma (RMS) cell lines

The human RMS cell line RD was a gift from Dr. Corinne Linardic. The RH30, RH36, Rh41, and Rh18 lines were obtained from Dr. Peter Houghton. All lines except RD were maintained in RPMI supplemented with 10% fetal bovine serum (VWR) at 37°C with 5% CO_2_. RD cells were maintained in DMEM supplemented with 10% FBS at 37°C with 5% CO_2_. Cell lines were authenticated by genotyping.

### Lentiviral/siRNA/retroviral knockdown assays

Scrambled-control and *SNAI2* -specific shRNAs were delivered via the pLKO.1-background vector and packaged using 293T cells. siSlug2 (SNAI2 sh1) was a gift from Bob Weinberg (Addgene plasmid # 10904; http://n2t.net/addgene:10904; RRID:Addgene_10904)^24^ siSlug3 (SNAI2 sh2) was a gift from Bob Weinberg (Addgene plasmid # 10905; http://n2t.net/addgene:10905; RRID:Addgene_10905)^24^. pMKO shRNA Bim was a gift from Joan Brugge (Addgene plasmid # 17235; http://n2t.net/addgene:17235; RRID:Addgene_17235)^34^. Retroviral particles were made in Plat-A packaging cells using TranstIT-LT1 (Mirus). RMS cells were infected with viral particles for 24 hr at 37°C using 8 mg/mL of polybrene (EMD Millipore).

### Cell confluence and colony formation assays

Rh18, Rh30, and RD parental cells were seeded into 24-well plates at 20% confluency and stored at 37°C in the Incucyte ZOOM (Essen Bioscience). After reaching ~40% confluency, cells were subjected to varying degrees of IR (0 Gy – 20 Gy) and placed back in the Incucyte. Total confluency over time was monitored every 4 hours over a period of 5 days. Incucyte confluency assays were performed similarly for scrambled, *SNAI2* shRNA, *TP53* shRNA, *BIM* shRNA, and *SNAI2/BIM* double knockdown shRNA RMS cells with and without exposure to IR. For colony formation assays, RMS cells were seeded in 12 well plates (~1,250 – 10,000 cells/well for cells receiving radiation and 300 – 600 cells/well for cell receiving no treatment). After 24h, cells were subjected to varying degrees of radiation (2 – 8 Gy). Incubation time for colony formation assays between cell lines varied from 3 to 6 weeks. When colonies were sufficiently large, media was gently removed from each plate by aspiration, and colonies were fixed with 50% methanol for 10 – 15 minutes RT. Colonies were then stained with 3% (w/v) crystal violet in 25% methanol for 10 – 15 minutes RT, and excess crystal violet was washed with dH_2_O with plates being allowed to dry. Colony formation was analyzed using ImageJ (Fiji). Significance was calculated by one-way ANOVA with Dunnett’s multiple comparisons tests. All assays were performed in triplicates.

### Western blot analysis

Total cell lysates from human RMS cell lines and human myoblasts were obtained following lysis in RIPA lysis buffer supplemented with protease inhibitors (Roche). Western blot analysis was performed similar to Ignatius et. al., 2017^49^. Membranes were developed using an ECL reagent (Western Lightning Plus ECL, PerkinElmer; or sensitive SuperSignal West Femto Maximum Sensitivity Substrate, Thermo Scientific). Membranes were stripped, rinsed, and re-probed with the respective internal control antibodies. List of primary and secondary antibodies is included in supplementary data (Supplemental Table 2).

### Immunohistochemistry

Once tumors reached 4X the initial volume mice were euthanized and tumors were fixed with 4% paraformaldehyde/PBS, sectioned, blocked, and stained with H&E, Ki67, Myogenin, and MF20 antibodies (see Supplemental Table 2).

### Caspase-Glo 3/7 assay

Rh18, Rh30, and RD Scr- and SNAI2-knockdown cells were seeded at 20% confluency in 24-well plates and placed in the Incucyte ZOOM instrument. After reaching ~40% confluency, media was supplemented with Caspase-Glo 3/7 reagent (1:1,000, Essen Bioscience) and Nuclight reagent (1:500, Essen Bioscience). Cells were then subjected to a range of doses of IR (0 Gy – 15 Gy) and placed back in the Incucyte. Images taken at 48h and 72h were processed using Adobe Photoshop and analyzed using ImageJ Cell Counter to determine percent caspase 3/7 events. Significance was determined using a two-way ANOVA with Sidak’s multiple comparisons test or one-way ANOVA with a post-hoc Tukey test accordingly.

### Flow Cytometry

Rh18, Rh30, and RD Scr- and SNAI2-knockdown cells were seeded in 6-well plates and irradiated (PXi Precision X-Ray X-RAD 320). Cells were collected at varying time points (48, 72, 96, and 120 hours). A negative control of cells that did not receive IR were collected as well. Cells were centrifuged and resuspended in annexin-binding buffer. After determining cell density and diluting to 1 x 10^6^ cells/mL with annexin-binding buffer, annexin V conjugate, and propidium iodide were added to sample aliquots and left to incubate at room temperature in the dark for 15 minutes. After incubation, aliquots were mixed gently while adding annexin-binding buffer on ice and analyzed by flow cytometry (LSRFortessa X-20; BD Biosciences). Cell cycle was assessed using the same cells and conditions described above with Click-iT EdU Alexa Flour 647 Flow Cytometry Assay (ThermoFisher) according to the provided protocol. Significance was determined using a Two-Proportion Z-test.

### RNA-seq

RNA was extracted using the RNeasy mini kit (Qiagen). Poly-A-selected RNA libraries were prepared and sequenced on an Illumina HiSeq2000. QC was performed using FastQC version 0.11.2 and Picard’s version 1.127 RNASeqMetrics function with the default parameters. PCR duplicates were marked using Picard’s MarkDuplicates function. RNA-seq reads were aligned to the UCSC hg19 reference genome using TopHat version 2.0.13. Significance was defined as having FDR *q*-value <0.01 and FWER *p*-value of <0.05. Gene set enrichment analysis (http://www.broadinstitute.org/gsea/index.jsp) was performed using default parameter settings.

### ChIP-seq

ChIP-seq data used, was published previously^30^ and performed as follows. 1% Formaldehyde-fixed chromatin from RD and SMS-CTR cells were sheared to 200-700 bp with Active Motif EpiShear Sonicator. Chromatin-IP with SNAI2 Ab (CST, Catalogue # 9585) was performed O/N, using ChIP-IT High Sensitivity kit (Active Motif). Drosophila chromatin (Active Motif, Catalogue #53083) and H2Av ab (Active Motif, Catalogue #61686) was used for spike-in normalization across samples. ChIP-seq libraries were prepared using Illumina TruSeq ChIP Library Prep Kit and sequenced on NextSeq500. Reads were mapped to reference genome (version hg19) using BWA. High-confidence ChIP-seq peaks were called by MACS2.1. Raw sequencing data and processed files are available through GEO (GSE137168).

### Gene expression analysis

Real-time qPCR was completed using the QuantStudio 7 Flex system (Applied Biosciences). PCR primers and specific conditions are provided in Table X and Supplemental Experimental Procedures. RNA isolation and cDNA preparation were performed as previously described^30,49^. Significance was calculated by a two-way ANOVA with a Sidak’s multiple comparisons test.

## Author contributions

LW, NH, KBo, performed and/or interpreted or supervised aspects of the different experiments and helped write the manuscript. BEG designed and implemented scripts, pipelines and analysis tools for RNA-seq and ChIP-seq. LW and MI initiated the experimental study. LW, NH, KBa, RMC, PS, JC, XZ helped with shRNA- and siRNA-mediated silencing experiments. EC, a soft tissue pathologist, helped with histology and immunohistochemistry analysis. KBo and LW performed all animal experiments. PH discussed the clinical importance of the results and the murine xenograft experiments. MI, with contributions from JK and PH, provided overall study direction, funding, supervision, and revised the manuscript. All authors critically reviewed the report and approved the final version.

---

**Supplemental Figure 1.**
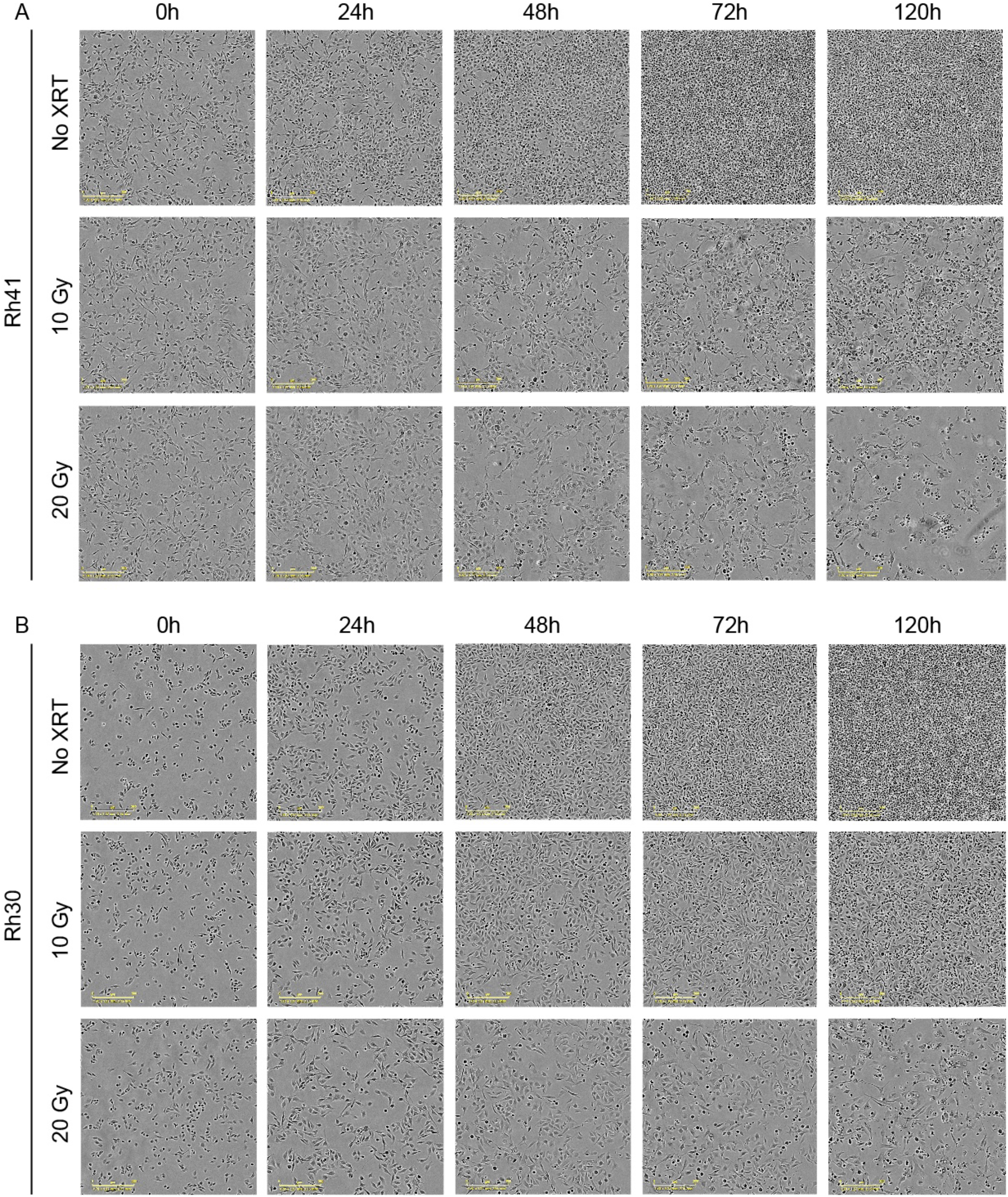
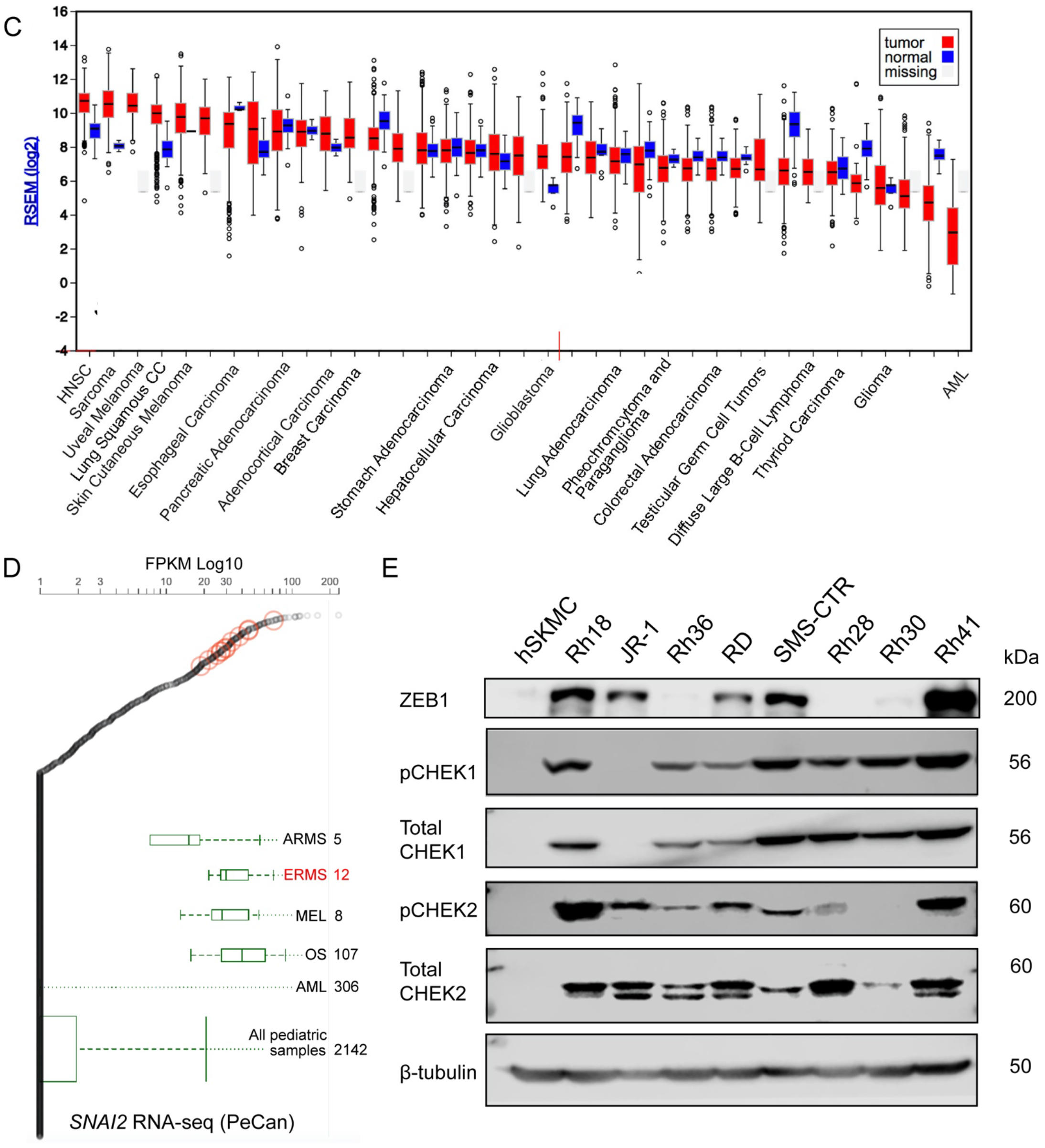
A, B. Growth of Rh41 and Rh30 cells after irradiation with increasing doses of IR at various time points post exposure. Scale bar 300 μm. C. *SNAI2* expression levels in tumors in the TCGA database. D. *SNAI2* expression in RMS tumors compared to other pediatric cancers, with SNAI2 expression levels in ERMS tumors circled in red (ARMS – alveolar rhabdomyosarcoma, ERMS – embryonal rhabdomyosarcoma, MEL – melanoma, OS – osteosarcoma, AML – acute myeloid leukemia; St Jude PeCan database)-Note this is the same data from Figure 1F but includes expression of *SNAI2* comparisons to other pediatric cancers. E. Western blots showing levels of ZEB, phosphorylated CHEK1 and total CHEK1, as well as phosphorylated CHEK2 and total CHEK2 across rhabdomyosarcoma cell lines; hSKMC, human Skeletal muscle cells were used as control for western blot analyses.

**Supplemental Figure 2.**
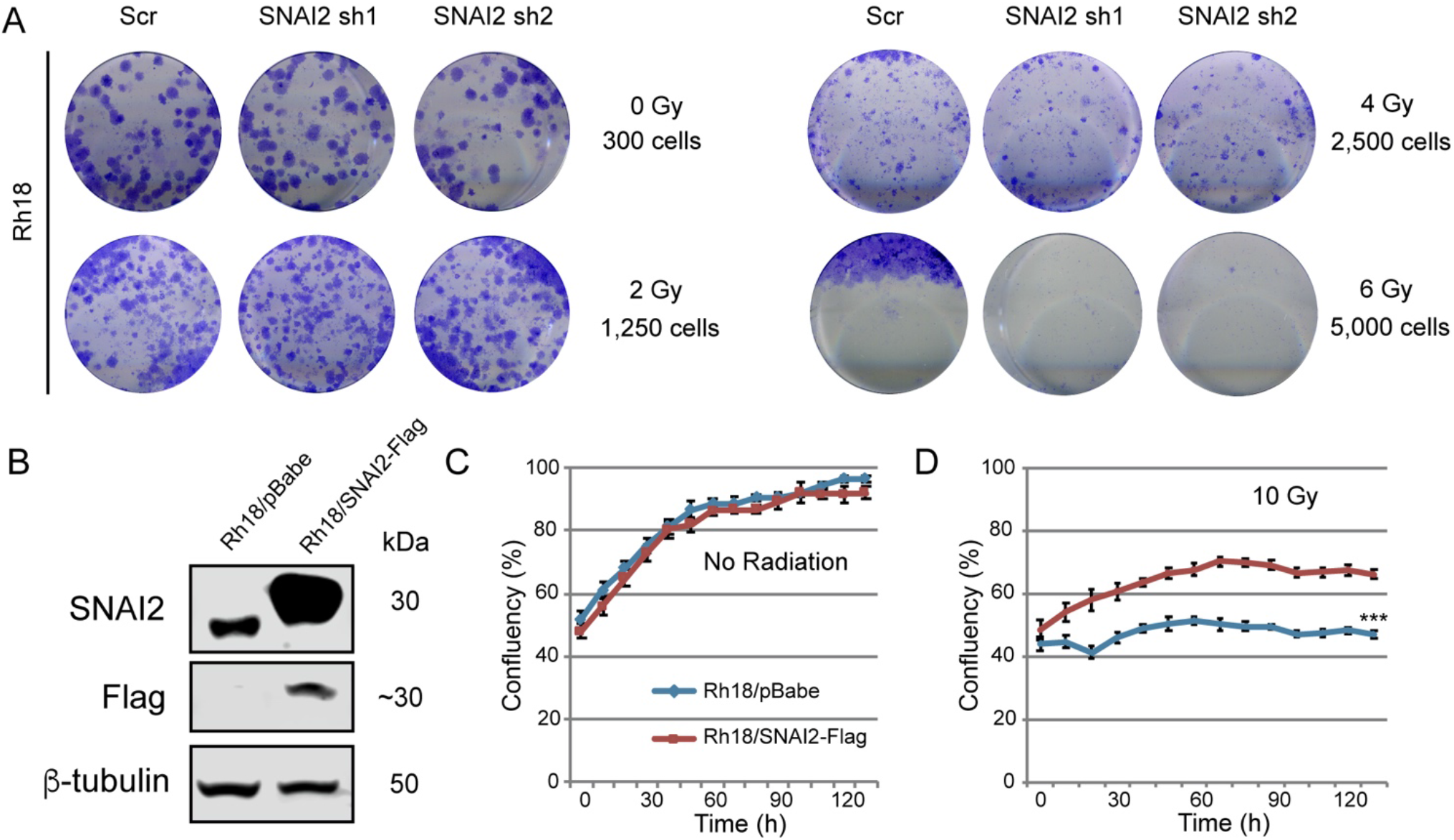
A. Representative colony formation wells for Rh18 Scr and SNAI2 knockdown (sh1 and sh2) after indicated IR exposure. B. Western blot of Rh18 pBabe control and SNAI2-Flag cells for SNAI2 and Flag expression. C. D. Confluency (%) of non-IR or IR-treated (10 Gy) Rh18 pBabe or SNAI2-Flag cells was assessed on phasecontrast images acquired from 0 to 130 h. ***p<0.001. Error bars represent ±1 SD.

**Supplemental Figure 3.**
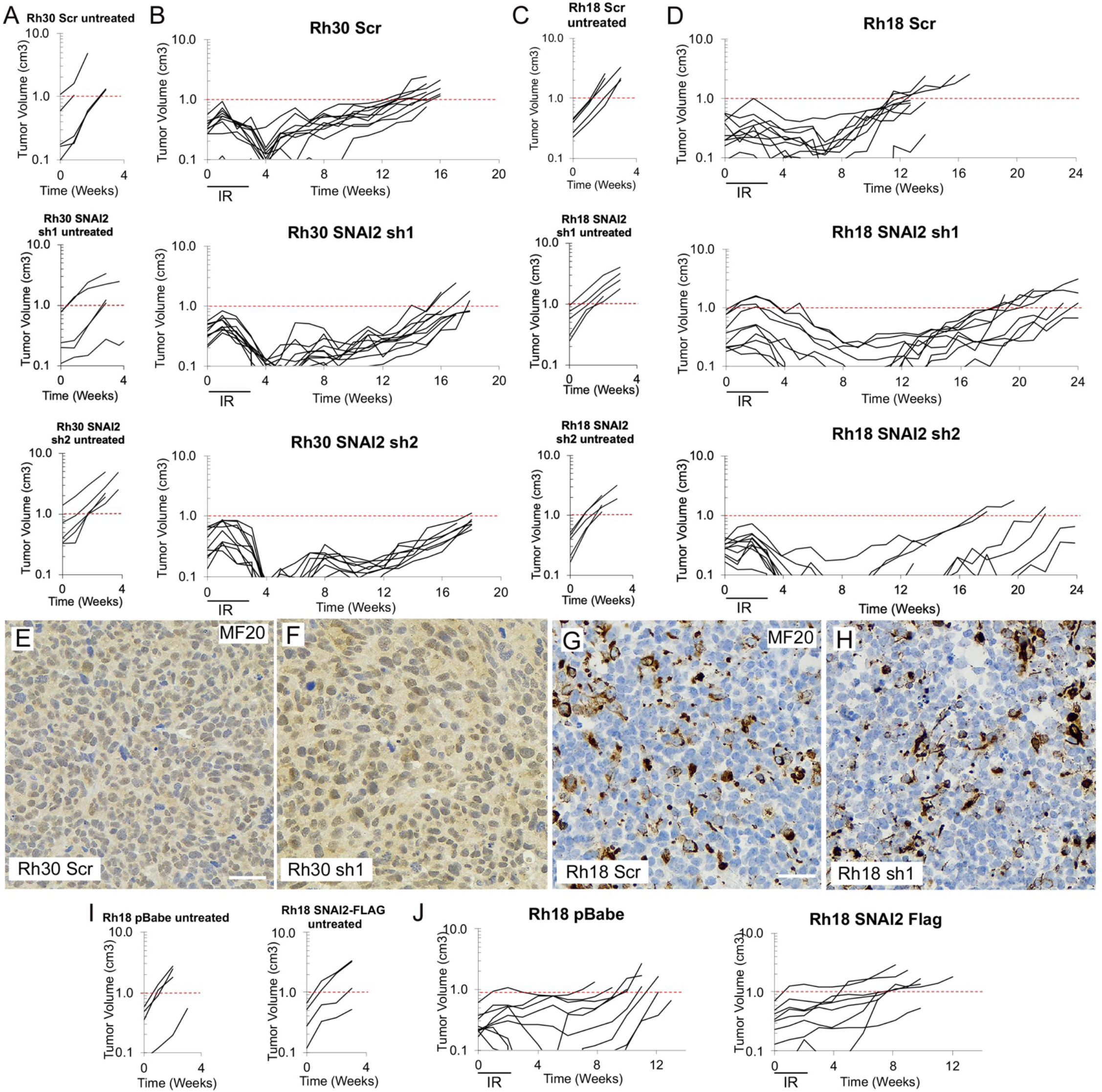
A. Individual tumor volumes for Rh30 Scr, SNAI2 sh1, and SNAI2 sh2 xenografts under non-IR conditions. B. Individual tumor volumes for Rh30 Scr, SNAI2 sh1, and SNAI2 sh2 xenografts after IR exposure (2 Gy/day, 5x a week, for 3 weeks). C. Individual tumor volumes for Rh18 Scr, SNAI2 sh1, and SNAI2 sh2 xenografts under non-IR conditions. D. Individual tumor volumes for Rh18 Scr, SNAI2 sh1, and SNAI2 sh2 xenografts after IR exposure (2 Gy/day, 5x a week, for 3 weeks). E, F. Representative immunohistochemistry sections of MyHC (MF20) staining in Rh30 Scr and SNAI2 sh1 xenografts. No statistical difference detected. Scale bar = 100 μm. G, H. Representative immunohistochemistry sections of MyHC (MF20) staining in Rh18 Scr and SNAI2 sh1 xenografts. No statistical difference detected. Scale bar = 100 μm. I. Individual tumor volumes for Rh18 pBabe and SNAI2-Flag xenografts under non-IR conditions. Red dashed line indicates 1 cm^3^ in all graphs. J. Individual tumor volumes for Rh18 pBabe and SNAI2-Flag xenografts after IR exposure (2 Gy/day, 5x a week, for 2 weeks).

**Supplemental Figure 4.**
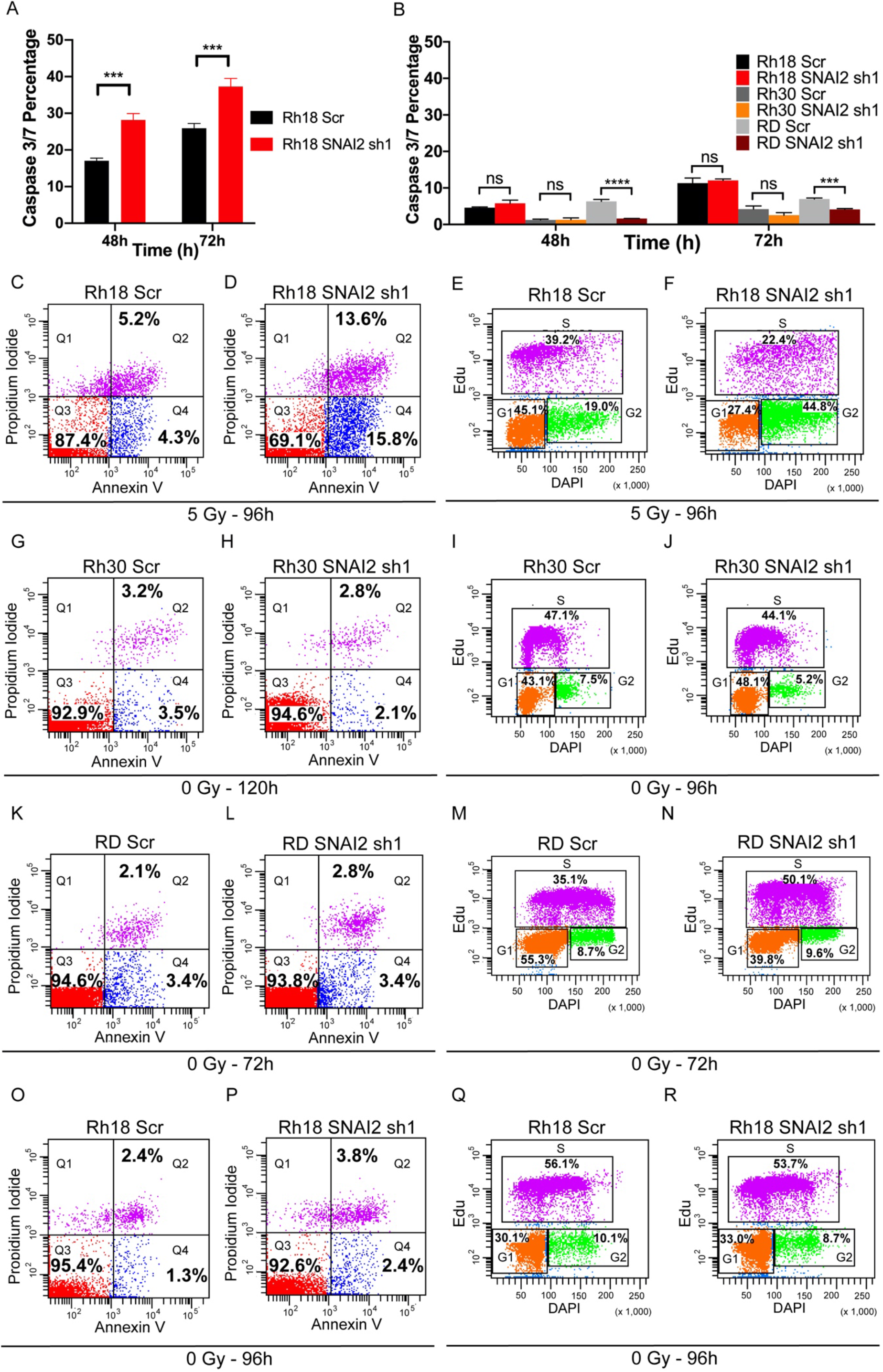
Related to Figure 4; Loss of SNAI2 promotes IR-mediated apoptosis and blocks irradiated RMS cells from exiting the cell cycle. A. Average Caspase 3/7 percentages (mean ± 1 SD) of Rh18 cells with either Scr shRNA or SNAI2 shRNA undergoing apoptosis after a 5 Gy IR dose. \*\*\**p*<0.001. B. Average Caspase 3/7 percentages (mean ± 1 SD) of Rh18, RD, and Rh30 cells (either Scr shRNA or SNAI2 shRNA) undergoing apoptosis grown under non-IR conditions. ns = not significant, \*\*\*\**p*<0.0001. C. D. Flowcytometry plots showing Propidium iodide vs. Annexin V staining in Rh18 cell lines with either Scr shRNA or SNAI2 shRNA knockdown after treatment with a 5 Gy IR dose. Rh18 early apoptosis (Q4): Scr 4.3% vs. sh1 15.8%, *p*<0.0001, late apoptosis (Q2): Scr 5.2% vs. sh1 13.6%, *p*<0.0001. E, F. Flowcytometry plots of EdU vs. DAPI staining in Rh18 cells with either Scr shRNA or SNAI2 sh1 after exposure to 5 Gy. Rh18 vs. sh1 G2 phase *p*<0.0001. G, H, K, L, O, P. Flow cytometry plots showing Propidium iodide vs. Annexin V staining in Rh18, RD, and Rh30 cell lines with either Scr shRNA or SNAI2 shRNA knockdown under growth conditions without IR. Not significantly different. I, J, M, N, Q, R. Flow cytometry plots showing cell cycle analysis with EdU vs. DAPI staining in Rh18, RD, and Rh30 cells with either Scr shRNA or SNAI2 shRNA under growth conditions without IR. Not significantly different.

**Supplemental Figure 5.**
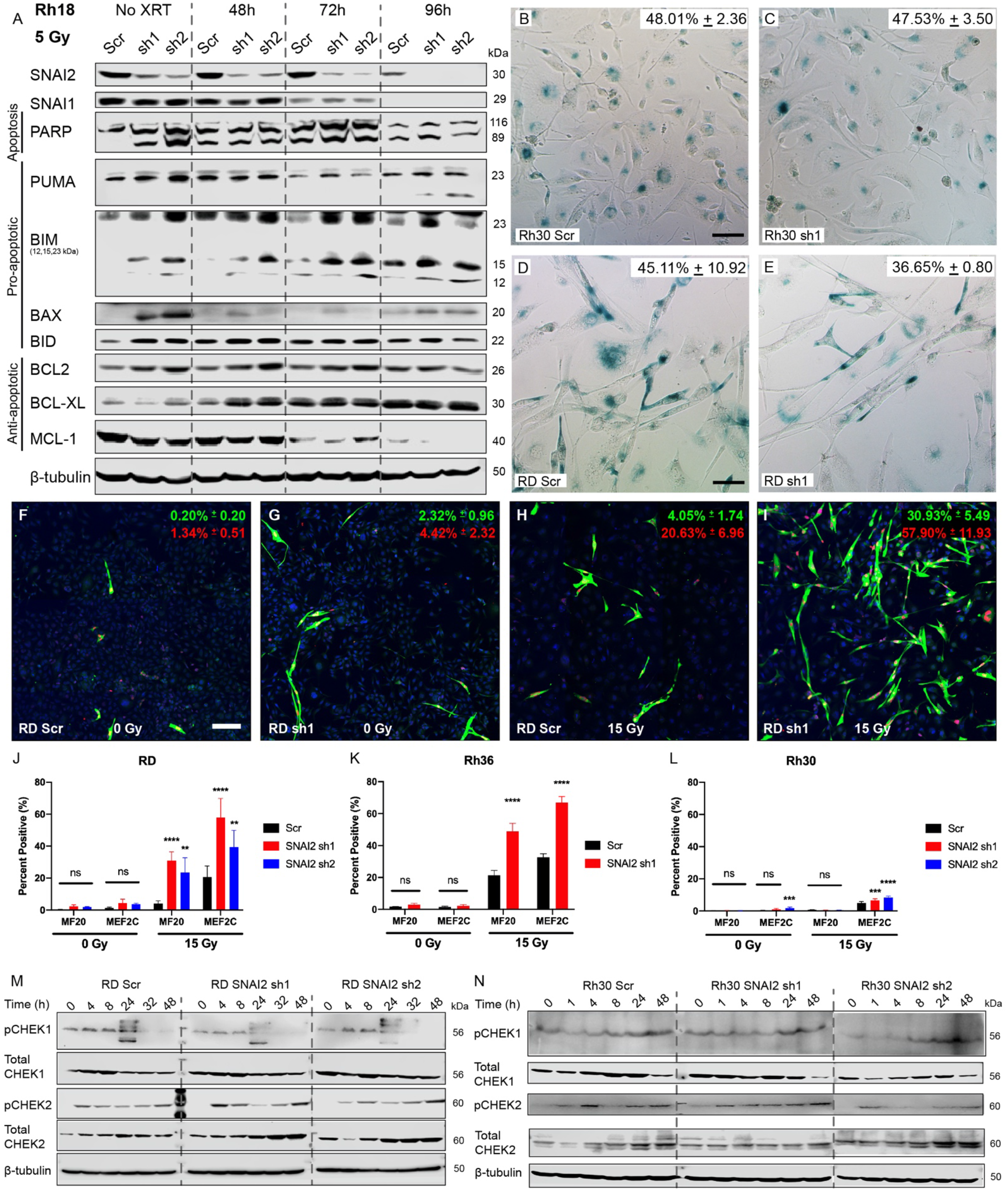
A. Western blot showing levels of apoptotic, pro-apoptotic, and anti-apoptotic proteins in Rh18 cells with control or SNAI2 knockdown, either non-irradiated or irradiated and assessed at 48, 72, and 96 hours post irradiation with 5 Gy. B-E. β-gal staining in Rh30 and RD cells with either Scr shRNA or SNAI2 shRNA treatments at 120h post IR (20 Gy). No statistical difference in percentage of β-gal staining. Scale bar = 5 μm F-I. Representative confocal microscopy images of RD cells with either Scr or SNAI2 shRNA expression immunostained with differentiated myosin MF20 and MEF2C antibodies under non-IR (F, G) or 15 Gy (H, I) IR conditions. Scale bar = 100 μm. J-L. Quantification of average MF20 and MEF2C in either non-IR or 15 Gy IR conditions in Scr vs SNAI2 shRNA treated RD, Rh36, and Rh30 cells. ns = not significant, \*\**p*<0.005, \*\*\**p*<0.001, \*\*\*\**p*<0.0001. Error bars represent ±1 SD. M. Western blots showing expression of phosphorylated CHEK1 and total CHEK1, as well as phosphorylated CHEK2 and total CHEK2, in RD cells with either Scr shRNA or SNAI2 shRNA treatments. N. Western blot showing levels of phosphorylated CHEK1 and total CHEK1 as well as phosphorylated CHEK2 and total CHEK2 in Rh30 cells with either Scr shRNA or SNAI2 shRNA treatments.

**Supplemental Figure 6.**
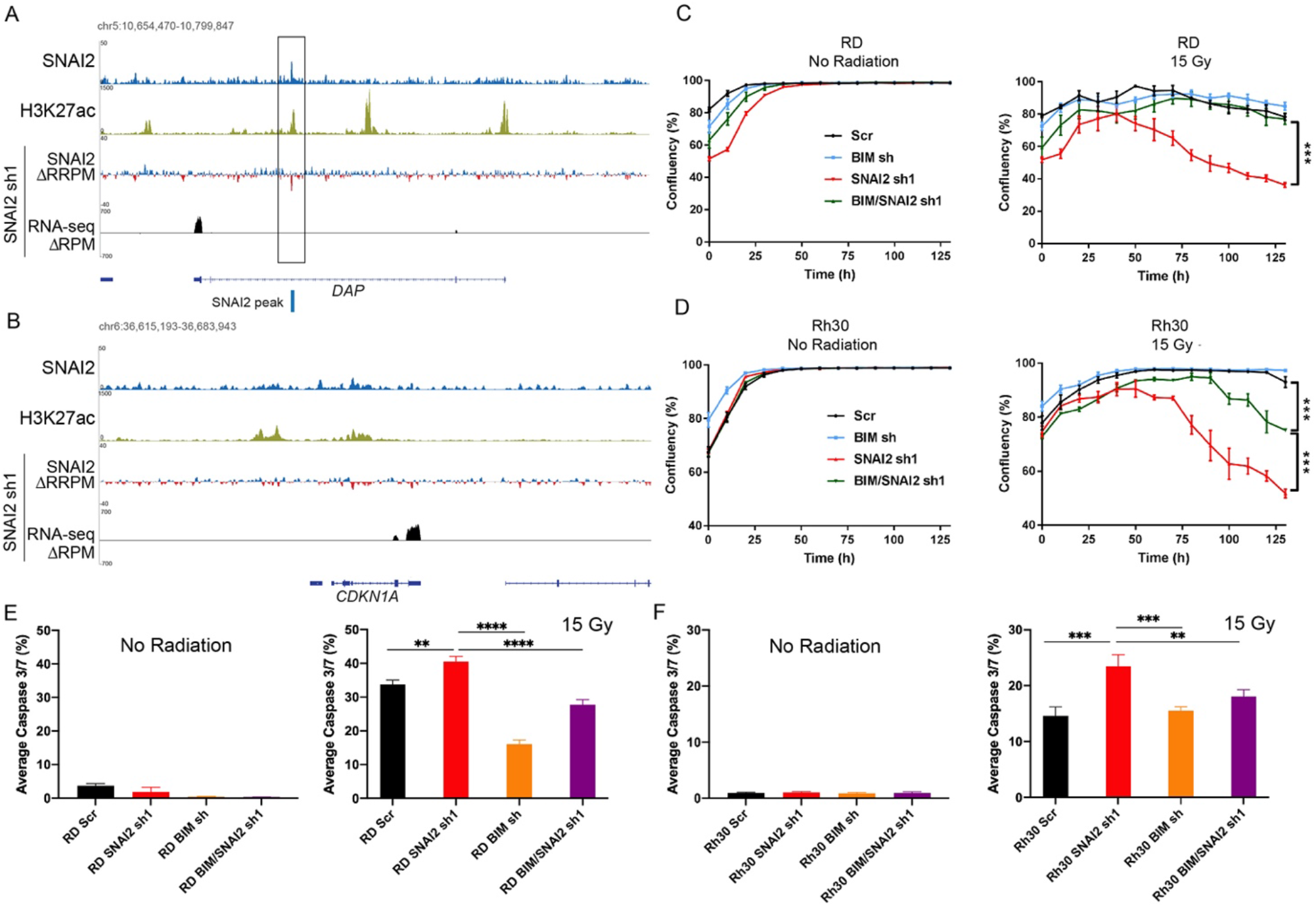
A, B. ChIP-seq tracks of SNAI2 (Blue) and H3K27ac (Yellow) binding in RD cells in control /Scr, delta (Δ) enrichment value (shSNAI2 sh1 minus shScr, Blue and Red) for SNAI2 and gene expression (RNA-seq, Black) for (A) *DAP* and (B) *CDKN1A.* Blue line represents SNAI2 called peak. Values on Y-axis represent fold enrichment. RRPM, Reference-adjusted Reads Per Million Mapped Reads; RPM, Reads Per Million Mapped Reads. C. Confluency (%) of non-IR or IR (15 Gy) treated RD cells expressing (either control, SNAI2, BIM, or BIM/SNAI2 shRNAs) was assessed using Incucyte Zoom software based on phase-contrast images acquired from 0 h to 125 h. \*\*\**p*<0.001. Error bars represent ±1 SD. D. Confluency (%) of non-IR or IR (15 Gy) Rh30 cells (either control, SNAI2, BIM, or BIM/SNAI2 knockdown) was assessed using Incucyte Zoom software based on phase-contrast images acquired from 0h to 125 h. \*\*\**p*<0.001. Error bars represent ±1 SD. E. Average Caspase 3/7 (%) (mean ± 1 SD) of RD Scr, SNAI2 sh1, BIM sh, BIM/SNAI2 sh1 cells under non-IR conditions at comparable densities as RD cells 72h after 15 Gy IR. 15 Gy values shown as reference (Figure 6F). No significant difference between cells in non-IR conditions. F. Average Caspase 3/7 (%) (mean ± 1 SD) of Rh30 Scr, SNAI2 sh1, BIM sh, BIM/SNAI2 sh1 cells under nonIR conditions at comparable densities as Rh30 cells 72h after 15 Gy IR. 15 Gy values shown as reference (Figure 6H). No significant differences between cells in non-IR conditions.

## Supplemental Materials and Methods

### TCGA Analysis/ PECAN expression analysis

Analysis of *SNAI2* expression across the TCGA database confirmed sarcoma tumors highly express *SNAI2* compared to other cancer types. Analysis of *SNAI2* expression in a different cohort of approximately 2000 pediatric cancers from the St. Jude-PeCan portal confirmed that *SNAI2* is highly expressed in RMS tumors and especially the ERMS sub-type compared to other pediatric cancers with osteosarcoma tumors expressing higher *SNAI2* (Figure 6).

### Senescence Cell Histochemical Staining

RMS cells were seeded into 6 well plates (0.1 − 0.5 x 10^6^ cells/well) and radiated after 24h. Cells were fixed after 120h with 4% paraformaldehyde. Cells were then stained using the Senescence Cells Histochemical Staining Kit (Sigma Aldrich) according to the provided protocol. Cells were allowed to incubate with the stain at 37°C without CO_2_ until cells were stained blue (2 hours to overnight). Percentage of cells positive for β-galactosidase was assessed using ImageJ. Significance was determined using Student’s t test.

### Immunofluorescence Staining

Immunofluorescence staining was performed similar to in Ignatius et. al., 2017^49^. Cells were plated at 4,000 cells/well (no IR) and 10,000 cells/well (receiving IR), grown in 10% FBS DMEM or RPMI growth media, fixed at 72 hpIR (0 or 15 Gy) in 4% paraformaldehyde/PBS, permeabilized in 0.5% Triton X-100/PBS, and incubated with rabbit anti-MEF2C (CST; Catalog No. 5030) and anti-myosin heavy chain (DSHB) in 1% BSA/PBS. Secondary antibody detection was performed with Alexa Flour 488 goat anti-mouse and Alexa Fluor 594 goat anti-rabbit (Invitrogen). Cells were counterstained with DAPI (1:10,000) and imaged. Images were processed in ImageJ and Adobe Photoshop.

